# Precancer: Mutant clones in normal epithelium outcompete and eliminate esophageal micro-tumors

**DOI:** 10.1101/2021.06.25.449880

**Authors:** B. Colom, A. Herms, M.W.J. Hall, S.C. Dentro, C. King, R.K. Sood, M.P. Alcolea, G. Piedrafita, D. Fernandez-Antoran, S.H. Ong, J.C. Fowler, K.T. Mahbubani, K. Saeb-Parsy, M. Gerstung, B.A. Hall, P.H. Jones

## Abstract

Human epithelial tissues accumulate cancer-driver mutations with age^1–7^, yet tumor formation remains rare. The positive selection of these mutations argues they alter the behavior and fitness of proliferating cells^8–10^. Hence, normal adult tissues become a patchwork of mutant clones competing for space and survival, with the fittest clones expanding by eliminating their less-competitive neighbors^9–12^. However, little is known about how such dynamic competition in normal epithelia impacts early tumorigenesis. Here we show that the majority of newly formed esophageal tumors are eliminated through competition with mutant clones in the surrounding normal epithelium. We followed the fate of microscopic tumors in a mouse model of esophageal carcinogenesis. Most neoplasms are rapidly lost despite no indication of tumor cell death, decreased proliferation, or an anti-tumor immune response. Deep-sequencing of 10-day and 1-year-old tumors shows evidence of genetic selection on the surviving neoplasms. Induction of highly competitive clones in transgenic mice increased tumor removal, while pharmacologically inhibiting clonal competition reduced tumor loss. The results are consistent with a model where survival of early neoplasms depends on their competitive fitness relative to that of mutant clones in the adjacent normal tissue. We have identified an unexpected anti-tumorigenic role for mutant clones in normal epithelium by purging early neoplasms through cell competition, thereby preserving tissue integrity.

## Main text

Somatic mutations in cancer-driver genes are frequent in phenotypically normal adult human epithelia such as that of the esophagus, skin, endometrium, lung and colon ^1–7^. Some of these mutations are more prevalent in normal epithelium than neoplastic lesions, leading to the suggestion that mutant clones in the non-neoplastic epithelium might have an unanticipated protective role against cancer^2,5^. Defining how mutant clones in normal epithelium and early neoplastic lesions compete for space and survival may be critical to understand the initial stages of tumorigenesis. However, despite the potential implications for cancer development, our knowledge of how mutant clones in normal tissues and tumors interact is limited.

Here we investigated this issue in the mouse esophageal epithelium (EE). This tissue consists of a single basal layer of progenitor cells that periodically divide and eventually stratify upwards forming a multi-layer of differentiated cells that are finally shed (**Extended Data Fig. 1a**). Dividing cells have an equal probability of generating progenitor and differentiating daughters, a balance that maintains tissue homeostasis (**Extended Data Fig. 1b and Supplementary information**) ^13,14^. Some mutations transiently tilt this balance towards higher production of progenitor cells, thus conferring a competitive advantage over their less fit neighbors (**Extended Data Fig. 1c**)^9,12^.

Administration of diethylnitrosamine (DEN), a mutagen present in tobacco smoke, in drinking water induces mutations in the epithelial cells of the mouse esophagus ^11^. Cells with competitive “winner” mutations form clones that expand laterally in the proliferative basal cell layer until they collide with other mutants of similar fitness, at which point they return towards neutral drift (**Extended Data Fig. 1d and Supplementary information**) ^11^. With time, mutant clones in the phenotypically normal mouse EE undergo genetic selection as they compete for the limited space (**Supplementary information**), and generate a mutational landscape similar to that found in the aging human normal esophagus ^11^. In addition, DEN treatment leads to the formation of discrete tumors scattered within this mutant patchwork ^9,15^, which makes this an ideal model to study interactions between mutant clones in normal and neoplastic epithelium.

### Most EE tumors are lost over time

1 year after DEN treatment mice exhibit an average of ~one tumor per EE ^15^. To study earlier steps in tumor development here we collected esophagi at different time-points, from as early as 10 days up to 18 months post-DEN treatment (**Fig. 1a**). After tissue collection the entire EE was 3D-imaged at single cell resolution by confocal microscopy, following immuno-staining of tumors with the stress marker protein KRT6 ^15^ (**Fig. 1b and Extended Data Fig. 2a-c**). This technique allowed us to detect hundreds of ‘micro-neoplasms’ per esophagus at 10 days post-DEN withdrawal. These were as small as ~50 μm in diameter and comprising as little as ~20 cells, with histological features of focal hyperplasia, (**Fig. 1b-d**). Remarkably, the number of tumors in the EE fell rapidly during the following months and stabilized after this initial phase (**Fig. 1d and Supplementary Table 1**). Persisting tumors kept growing and developed features of dysplasia, with angiogenesis in the underlying submucosa and leukocyte infiltration (**Fig. 1b, c, e; Extended Data Fig. 2c-e and Supplementary Table 1**). We conclude that most of the early EE tumors generated by DEN treatment are resolved soon after they are formed (**Fig. 1f**).

**Fig. 1.**
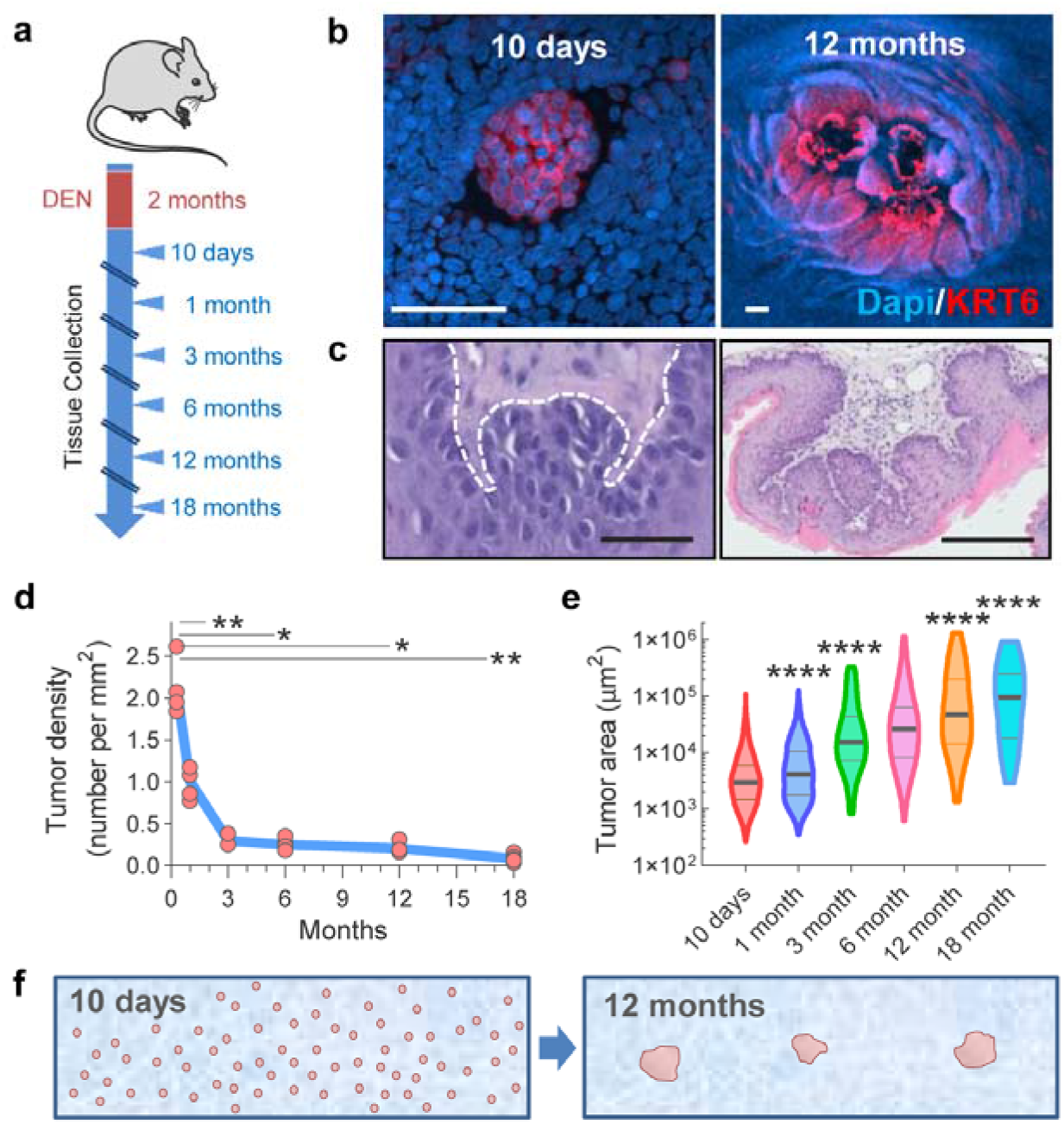
Most esophageal tumors are lost. (**a**) Wild-type mice received DEN in drinking water. The esophagus was collected at the indicated time-points. (**b-c**) Confocal (**b**) and H&E (**c**) images of EE tumors from mice collected 10-days or 12-months post-DEN treatment. Scale-bars: 50μm. Tissues in (**b**) were stained for Dapi (blue) and Keratin 6 (KRT6, red) to identify tumors. White-dots in (**c**) delineate the basal cell layer. (**d-e**) Density (**d**) and size (**e**) of tumors at the indicated time-points post-DEN treatment. Blue line in (**d**) connects the Mean values (Two-sided Mann-Whitney test, vs 10-days). Violin plot lines in (**e**) show median and quartiles (Two-sided Mann-Whitney test, vs 10-days). Number of mice (and tumors) are: 10d = 4(588), 1m = 5(408), 3m = 3(73), 6m = 5(92), 12m = 4(58) and 18m = 6(42). (**f**) Cartoon illustrating tissue tumor clearance over time. * p<0.05, ** p<0.01, **** p<0.0001.

### Sequencing of newly formed and surviving tumors

To gain insight into how this process may occur we next characterized and compared the mutational landscapes of newly formed and surviving tumors, collected at 10 days and 1 year post-DEN treatment, respectively (**Fig. 2a; Extended Data Figs. 3 and 4a**). Deep-targeted sequencing, with a median on-target coverage 236x, of 192 genes implicated in epithelial carcinogenesis identified a total of 11,149 mutations from 141 tumors (**Fig. 2b, Extended Data Fig. 4b and Supplementary Tables 2 and 3**). The mutational spectrum comprised mainly T>A/A>T, T>C/A>G and C>T/G>A alterations, consistent with DEN mutagenesis in mouse EE (Extended Data Fig. 4c) ^11^. Functionally, most mutations were protein altering, with missense SNVs being the most common, a pattern that was maintained over time (**Extended Data Fig. 4d-e**). To test whether genetic selection was a feature of tumor survival we calculated the dN/dS (non-synonymous-to synonymous) mutation ratios across all sequenced genes, with dN/dS values over 1 indicating positive selection ^2,3,16^. Mutant *Notch1* and *Trp53* were positively selected in 10 day neoplasms, whereas *Atp2a2, Notch1, Notch2, Chuk* and *Adam10* were selected in the surviving tumors at 1 year (**Fig. 2c and Supplementary Tables 4 and 5**). No additional driver genes were identified by dN/dS analysis of whole exome sequenced tumors (**Extended Data Fig. 5; Supplementary Tables 6-7 and Supplementary information**). Mutations in *Atp2a2* and Notch1 dominated clonal selection within tumors, exhibiting the highest gain in dN/dS values, number of mutations and estimated proportion of tumor area occupied by the mutant gene (**Fig. 2 c-d and Extended Data Fig. 4f**). Strikingly, over 98% of the persisting tumors at 1 year carried non-synonymous mutations in one or both of these genes (**Extended Data Fig. 4g**) Of note, whole exome and whole genome sequencing analysis showed that chromosomal alterations were negligible in these tumors, indicating that genome instability does not underpin tumor survival in our model (**Extended Data Fig. 6 and Supplementary information**). Together these results argue that the early generated EE neoplasms undergo genetic selection, a process that may influence their survival or elimination from the tissue.

**Fig. 2.**
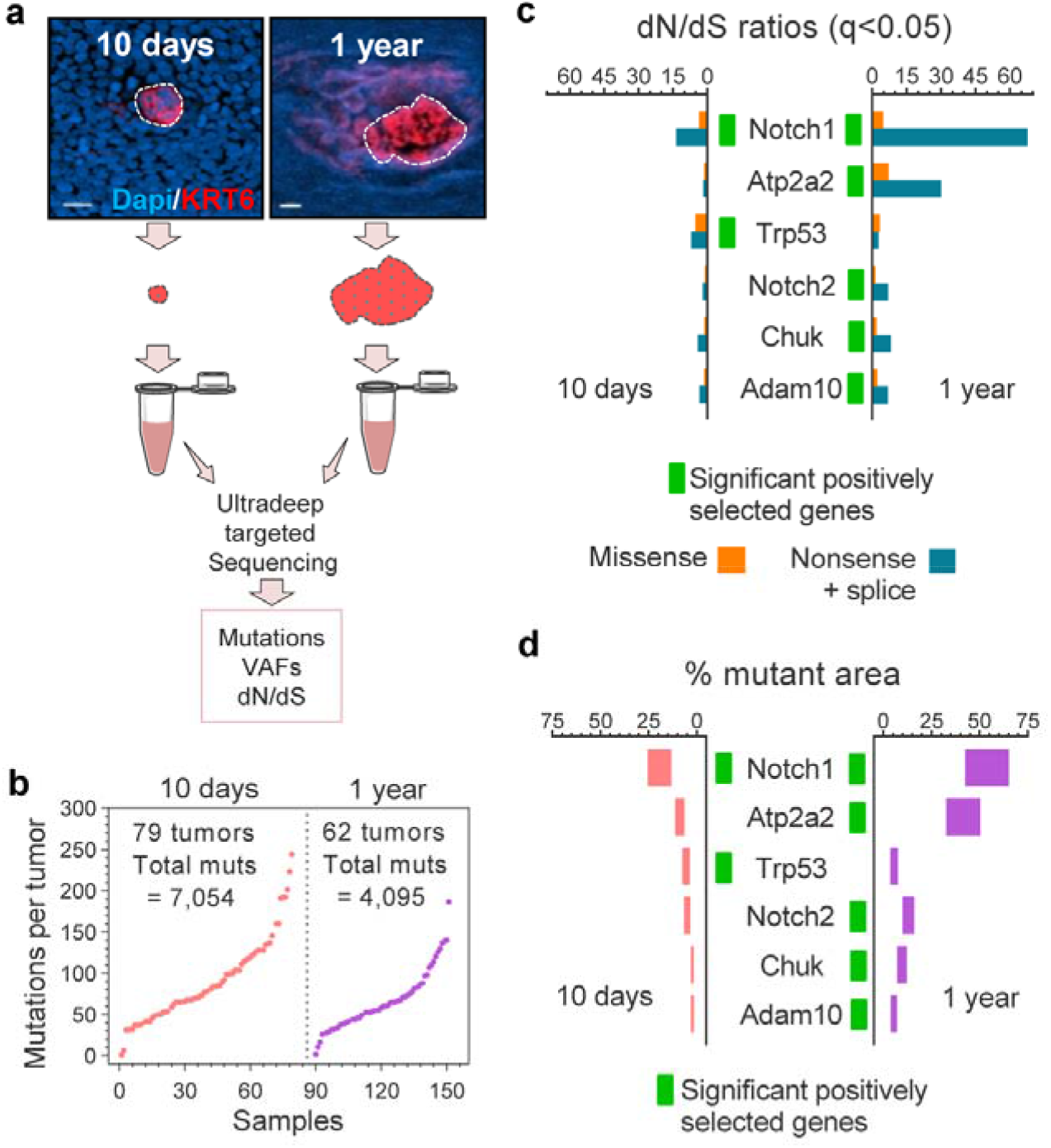
Sequencing of newly born and surviving tumors reveals patterns of selection. (**a**) EE tumors were collected at 10-days and 12-months post-DEN treatment as in **Figure 1**, and ultra-deep targeted sequenced using a 192 gene bait-set. Scale-bars: 50μm. (**b**) Number of mutations per tumor, including essential splice, frameshift, missense, nonsense and silent mutations. (**c**) dN/dS ratios for missense and truncating (nonsense + essential splice site) substitutions indicating genes under significant positive selection in the tumors (green rectangles, q<0.05 calculated with R package dNdScv ^16^). (**d**) Estimated percentage of tumor area carrying non-synonymous mutations for each gene. Range indicates upper and lower bound estimates. Green rectangles indicate dN/dS positively selected genes for each time-point.

### Mechanisms of early tumor loss

We next sought to identify the mechanism(s) responsible for early tumor loss. Immuno-staining of activated caspase 3, EdU (5-ethynyl-2⍰-deoxyuridine) short-term DNA incorporation assays and label-retaining experiments using *Rosa26^M2rtTA^/TetO-HGFP* mice, demonstrated that early tumors were not lost by apoptosis or abnormal proliferation of tumor cells (**Fig. 3a-d, Extended Data Fig. 7a-c, Supplementary Table 8 and Supplementary information**). We next explored whether the immune system could be eliminating these early neoplasms. However, although immune (CD45^+^) cells were occasionally observed in close contact with early tumors, the density of immune cells within them was no higher than in the surrounding tissue (**Extended Data Fig. 7d-j and Supplementary information**). Furthermore, immune-deficient *NOD.Cg-Prkdc^scid^ Il2rg^tm1Wjl^*/SzJ mice, which lack the typical anti-tumor immune response, showed equivalent tumor formation and loss rates as compared to wild-type controls. These results argue that the immune system is neither implicated in the initiation or elimination of early tumors, and that tumor removal takes place prior these being detected by the immune system (**Fig. 3e-f and Supplementary information**).

**Fig. 3.**
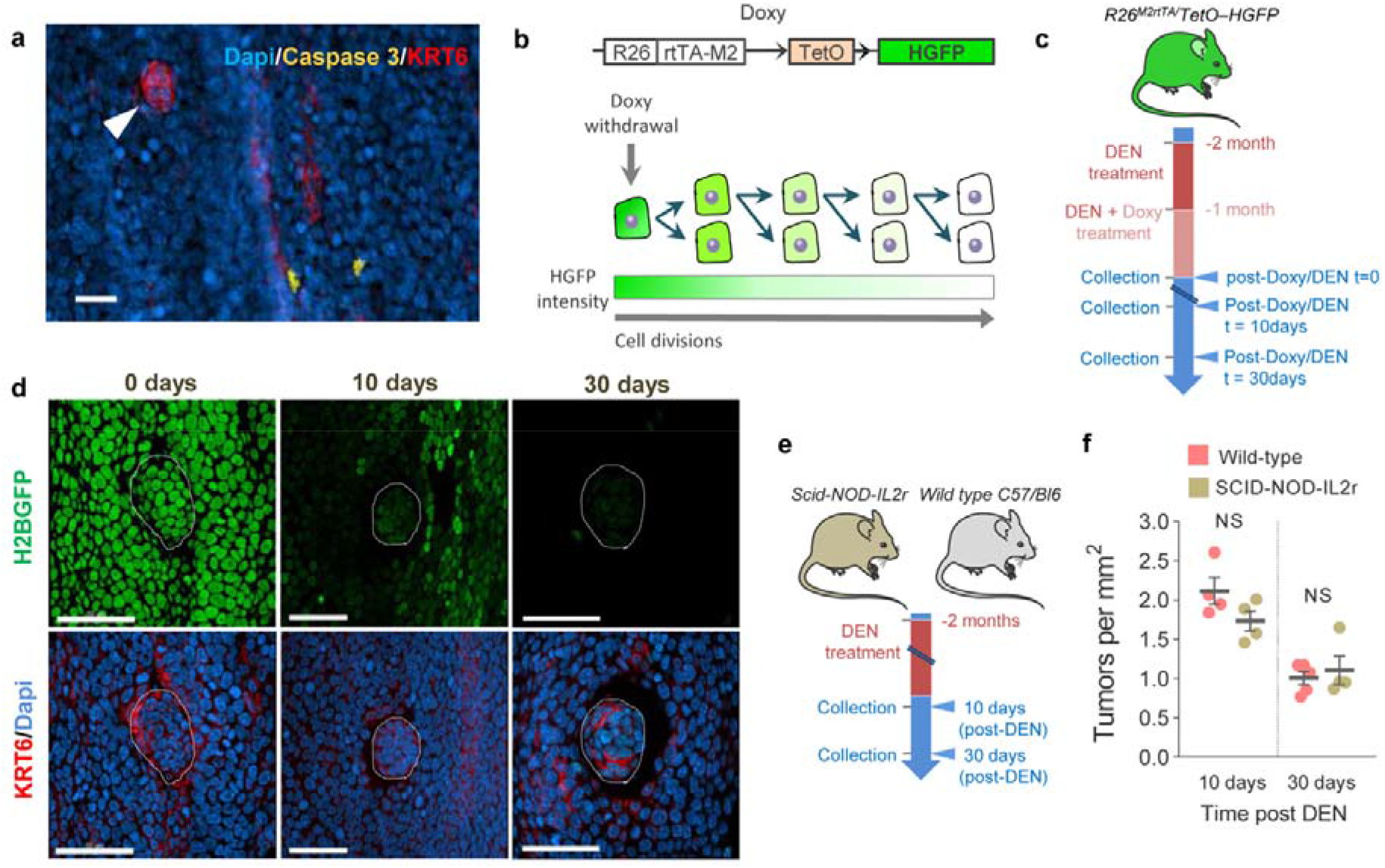
Early tumors are not eliminated by tumor cell apoptosis, abnormal proliferation or the immune system. (**a**) Representative confocal image of a 10-day post-DEN EE immuno-stained for activated Caspase 3^+^ and KRT6. The image shows two apoptotic cells in the normal epithelium (yellow). No apoptotic cells were detected in the tumors (arrowhead) (0/23 tumors analyzed from 2 mice). Scale-bar: 20μm. (**b**) *In vivo* label-retaining assay using **Rosa26^M2rtTA^/TetO-HGFP** transgenic mice to measure the rate of progenitor cell division. In this strain, expression of a stable Histone-Green Fluorescent Protein (HGFP) is activated by treatment with Doxycycline (Doxy). When Doxy is withdrawn, levels of the stable protein fall at a rate dependent on the rate of cell division. (**c**) Protocol: **Rosa26^M2rtTA^/TetO-HGFP** mice received DEN for one month followed by DEN + Doxy for another month. Tissues were collected at times 0, 10 and 30 days after Doxy/DEN withdrawal. (**d**) Confocal images showing HGFP expression in tumors (dotted lines) and their normal adjacent EE at different time-points post-Doxy withdrawal. Scale bars: 50μm. (**e**) Protocol: Wild-type and immunocompromised *Scid-NOD-IL2r* mice were treated with DEN for two months and the tissues collected 10 or 30 days after DEN withdrawal. (**f**) Number of tumors per mm^2^ of EE in wild-type and Scid-NOD-IL2r mice, collected 10-days (n=4) or 30-days (n=5 and 4, respectively) post-DEN treatment. Mean ± s.e.m. NS= not significant (Two-sided Mann-Whitney test).

### Mutant clones in normal EE remove tumors

We next investigated whether competition with mutant clones in the surrounding normal EE may act as a selective pressure for early tumor survival. We have previously described how highly competitive mutant clones in the normal EE of DEN-treated mice eliminate their less-fit neighbors (**Supplementary information**) ^11^. Thus, we hypothesized that a similar “survival of the fittest” competition mechanism could be responsible for the loss of early tumors (**Fig. 4a and Supplementary information**). To test this hypothesis, we analyzed whether induction of highly fit mutant clones in the normal epithelium altered early tumor survival (**Extended Data Fig. 8a-b and Supplementary information**). For this we used *Ahcre^ERT^Rosa26^wt/DNM-GFP^* (MAML-Cre) mice, which carry a highly competitive inducible dominant negative mutant allele of *Maml-1* (DN-Maml1) that inhibits Notch signaling (**Extended Data Fig. 8c**) ^9^. Induction of MAML-Cre mice 10 days post-DEN treatment accelerated tumor removal, proportionally to the expansion of DN-Maml1 mutant clones (**Fig. 4b-d and Extended Data Fig. 8d-g**). Analysis of images of tumors surrounded by DN-Maml1 clones suggest a mechanism of extrusion by competition, similar to that described in the skin and intestinal epithelium (**Fig. 4e and Extended Data Fig. 9a-c**) ^17–19^. The loss of tumors in these experiments cannot be explained by a model that does not include the effects of the surrounding mutant clones (**Supplementary information**). Instead, the results are compatible with a model where tumors are eliminated by competition with highly fit clones in the adjacent normal epithelium (**Supplementary information**). To challenge this hypothesis we explored the effect of neutralizing clonal competition following tumor formation. We have previously shown that Notch1 inactivating mutations confer the strongest competitive advantage in the DEN-treated EE ^11^. Therefore, inactivating Notch signaling by treating mice with Dibenzazepine (DBZ) would be expected to ‘level up’ the competitive fitness across the tissue and decrease tumor loss, as mutant epithelial clones would have no advantage over tumors ^9^ (**Extended Data Fig. 10a-b and Supplementary information**). The results show that DBZ treatment does indeed decrease tumor loss (**Extended Data Fig. 10c-d**). We conclude that expansion of mutant clones in the normal EE removes early neoplasms through competition, acting as a selective pressure for tumor survival (**Supplementary information**).

**Fig. 4.**
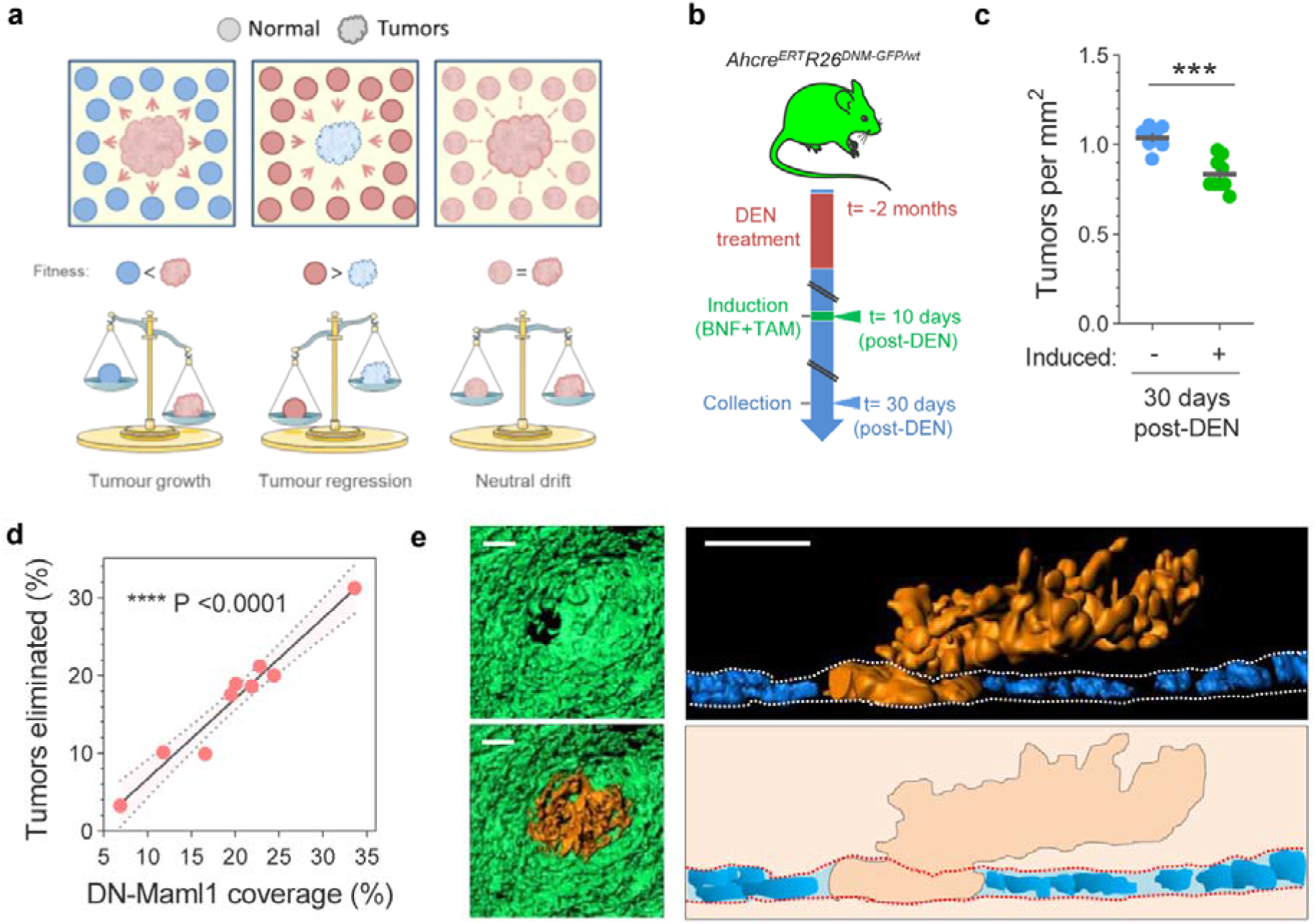
Elimination of early tumors by expansion of mutant clones in the adjacent normal epithelium. (**a**) In our model the survival of tumors is dependent on their relative competitive fitness compared to the surrounding normal tissue. This predicts 3 possible scenarios: 1) tumor cells are fitter than those in the normal adjacent epithelium (left), resulting in tumor growth; 2) Tumor cells are less fit than the adjacent normal (middle), resulting in tumor regression and elimination, and 3) cells in tumor and adjacent normal epithelium are equally fit (right), resulting in neutral competition. (b) Protocol: *Ahcre^ERT^Rosa26^wt/DNM-GFP^* mice (**Extended Data Fig. 10b**) received DEN for two months. 10 days after DEN withdrawal DN-Maml1 clones were induced and tissues harvested twenty days later. Control mice received DEN but were not induced. (**c**) Number of tumors per mm^2^ of EE in *Ahcre^ERT^Rosa26^wt/DNM-GFP^* induced and non-induced controls, as in (**b**). (**d**) Correlation between the area covered by DN-Mam1 clones and tumors being eliminated from the EE. Each dot represents a mouse (n=9). Line shows the Pearson correlation (R^2^=0.9541) with 95% confidence interval (colored area). (**e**) Top view (left), lateral view (right top) and schematic (right bottom) of a tumor (orange) surrounded by a DN-Maml1 clone (green). Dapi is shown in blue. Scale-bars: 20μm.

## Discussion

Our bodies are protected with highly efficient innate mechanisms to suppress tumors, typically exemplified by the immune system. In this study, we have identified a tumor-protective mechanism, independent of the immune system, by which expanding highly fit mutant clones present in normal EE outcompete and eliminate early neoplastic lesions. Importantly, the results imply that the survival of early neoplasms does not depend solely on the mutations they carry, but also on the mutational landscape of the neighboring normal tissue. These findings help reconcile the seemingly conflicting observations of a relatively low cancer incidence despite the large burden of cancer-associated mutations observed in healthy tissues ^1–7^, and suggest that some mutations found in normal tissues may contribute to the maintenance of homeostasis, with clear implications in cancer and aging.

Whether an analogous anti-tumor mechanism exists in humans is unknown. However, the existence of small neoplasms phenotypically similar to those found in our mutagenized mouse model, in otherwise healthy human EE (**Extended Data Fig. 10e-f**), together with the observed preferential colonization of Notch1-mutant clones in human normal EE versus squamous neoplasms ^2,5^, suggest the anti-tumor mechanisms described here may be common across species. Likewise, the high burden of cancer-driver somatic mutations identified in other normal human epithelia including that from skin, endometrium, lung and colon, suggests a similar mechanism of tumor purging by competition with mutant clones in adjacent normal epithelium could also take place in these tissues ^1–7^.

Our results show that modifying the relative competitive fitness of clones in normal epithelium and neoplastic lesions influences early tumor dynamics. Future studies are needed to explore whether environmental factors that alter the mutational landscape of normal tissues also impact early tumorigenesis through the mechanism described here. This study further suggests that treatments aimed at manipulating the mutant clonal architecture of normal epithelium might be exploited as an early cancer prevention strategy ^10,12^.

## Materials and Methods

### Mice

All animal experiments were conducted at the MRC ARES facility, Babraham or the RSF facility at the Wellcome Sanger Institute according to the UK Home Office Project Licenses 70/7543, P14FED054 or PF4639B40. Mice were housed in individually ventilated cages and fed on standard chow. Male and female adult mice were used for experiments. C57BL/6N wild-type mice and *NOD.Cg-Prkdc^scid^ Il2rg^tm1Wjl^/SzJ* ^22^ mice, which lack typical anti-tumor immune responses including a complete absence of B and T lymphocytes, NK cells, and defective dendritic cells and macrophages, were obtained from core colonies bred at the RSF facility. Double mutant *R26^M2rtTA^/TetO–HGFP* and *AhCre^ERT^Rosa26^flDNM−GFP/wt^* strains on a C57BL/6N background were generated as described previously ^9,13,15^.

### Chemical model of mouse esophageal neoplasia

Adult (>10 week old) mice were treated with Diethylnitrosamine (DEN) in sweetened drinking water (40⍰mg per 1,000⍰ml) for twenty-four hours, three days a week (Monday, Wednesday and Fridays) for eight weeks ^9,15^. Animals received sweetened water in between DEN dosages and normal water after DEN treatment was completed. Tissue (esophagus) samples were collected from 10 days to 18 months after DEN treatment.

### Mouse whole-esophagus sampling and processing

To prepare the esophagus for whole-tissue staining and imaging, tissues were dissected and placed in cold PBS. The esophageal tube was cut longitudinally and the muscle layer removed by pulling gently with micro-forceps. To separate the epithelium from the underlying submucosa the entire tissue was incubated for 2–3 hours in 5 mM EDTA at 37 °C and both layers were pulled away under a dissecting microscope with fine forceps. The whole esophageal epithelium was then flattened, fixed in 4% paraformaldehyde for 30 min at room temperature, washed in PBS and finally stored at 4 °C until further processing.

### Mouse whole-esophagus immunostaining

For immunostaining, whole-tissue esophageal epitheliums were blocked for one hour in 800 μl of staining buffer (0.5% bovine serum albumin, 0.25% fish skin gelatin, 0.5% Triton X-100 in PBS and 10% donkey serum) followed by an overnight incubation at room temperature with primary antibodies against KRT6, GFP, CD31, CD45 or Caspase 3, in staining buffer. Samples were then washed four times (x20 minutes) with 0.2% Tween-20 in PBS and, when necessary, incubated with secondary antibodies in staining buffer for three hours at room temperature. Samples were finally washed as above, incubated overnight at room temperature with 1 μg/ml DAPI or 0.4 μM TO-PRO™-3 Iodide solution to stain cell nuclei, and mounted using VECTASHIELD Mounting Media before image acquisition.

### Human samples

Esophagi were obtained from deceased organ donors. Informed consent for the use of tissue was obtained from the donor’s family (REC reference: 15/EE/0152 NRES Committee East of England - Cambridge South). A full thickness segment of mid-esophagus was excised within 60 minutes of circulatory arrest and preserved in University of Wisconsin (UW) organ preservation solution (Belzer UW^®^ Cold Storage Solution, Bridge to Life, USA) at 4 °C until further processing.

### Immunostaining of human esophagus

Donor esophageal samples were cut longitudinally, flattened and the muscle layer removed. Samples were blocked for one hour in 1mL of staining buffer (0.5% bovine serum albumin, 0.25% fish skin gelatin, 1% Triton X-100 and 10% donkey serum in PHEM buffer (60 mM PIPES, 25 mM HEPES, 10 mM EGTA, and 4mM MgSO4·7H20)), followed by two days incubation at room temperature with primary antibody against KRT6 in staining buffer. Samples were then washed for 2 days in PHEM buffer before incubation with secondary antibody in staining buffer for two days at room temperature. Samples were washed as above and incubated overnight at room temperature with 0.4 μM TO-PRO™-3 Iodide solution to stain cell nuclei. Finally, samples were incubated for one hour at 37 °C and then mounted, both in RapiClear 1.52 (SUNJin lab) solution, for image acquisition.

### Confocal microscopy

To obtain high quality confocal images of the entire mouse esophagus, whole-mounted tissues were imaged in a Leica TCS SP8 confocal microscope coupled to a high precision motorized stage. Contiguous 3D images of all epithelial layers (basal + suprabasal) were obtained and merged computationally using the mosaic function of the Leica Software. Typical settings for acquisition of multiple z stacks were 2.5μm z-step size, zoom x1, optimal pinhole, line average 4, scan speed 400⍰Hz and a resolution of 1,024 × 1,024 pixels using a 10X HC PL Apo CS Dry objective with a 0.4NA. Visualization and image analysis was performed using IMARIS (bitplane), ImageJ or Volocity 3D Image Analysis Software (Perkin Elmer). Otherwise, close up images were obtained with ×10, ×20 or ×40 objectives with typical settings for acquisition of z stacks being optimal pinhole, line average 3–4, scan speed 400-600⍰Hz and a resolution of 512 x 512 or 1,024 × 1,024 pixels.

### KRT6 staining as an early marker of mouse esophageal epithelial tumors

KRT6 is a keratinocyte stress-induced protein upregulated at both RNA and protein levels in mouse esophageal tumors ^15^. Here we used whole-tissue KRT6 staining to identify DEN-generated neoplasms in the esophageal epithelium from as early as 10 days post-DEN treatment. To characterize the protein expression levels in the tumors relative to the surrounding normal epithelium, the KRT6 staining profile in and around tumors was analyzed from esophagi collected 10 days after DEN treatment. The intensity of KRT6 and the nuclear marker DAPI were measured with ImageJ from defined ROIs containing tumors or their normal surrounding epithelia.

### Tissue histology

Esophagus from DEN-treated mice were dissected, fixed in 10% formalin for at least 24 hours and stored at 4°C. Tissues were then embedded in paraffin and cut at 5 μm thickness. Sections were stained with hematoxylin and eosin and scanned.

### Quantification of tumor number and size

To quantify the number and the area of the neoplasms present in the esophagus of carcinogen-treated mice at different time points, whole-tissues were stained with antibodies against KRT6 and a nuclear marker such as DAPI or TO-PRO™-3. High definition 3D confocal images of the whole esophagus were obtained and visualized with IMARIS software and ImageJ. Tumors were manually identified by shape (based on the nuclear staining) and KRT6 staining profile. The number of tumors quantified in the entire esophagus was divided by the total area of the tissue to obtain the number of tumors per mm^2^ of esophageal epithelium. The size of individual neoplasms was calculated by assuming an ellipse shape using the following formulae: A = (D_1_ D_2_ π) / 2, where D_1_ and D_2_ are the length of the major and minor diameters measured with IMARIS software.

#### Targeted sequencing of 10 days and 1 year esophageal neoplasms

##### Tissue preparation

Mice were treated with DEN for eight weeks and culled 10 days or 1 year after treatment. Mouse esophagi were dissected, cut longitudinally and the muscle layer removed. Tissues were incubated for 2–3 hours in 5 mM EDTA at 37 °C to separate the epithelium from the underlying submucosa. The whole epithelium was flattened, fixed in 4% paraformaldehyde for 30 min at room temperature and washed in PBS. Tissues were then immuno-stained for KRT6 and the nuclei labelled with DAPI or TO-PRO™-3 to identify the tumors before being imaged by confocal microscopy as described above.

##### Excision and digestion of neoplasms

Esophageal neoplasms from animals culled 10 days or 1 year post-DEN treatment were identified from the confocal images and manually cut using micro-scalpels under a fluorescent dissecting microscope. Tumors were placed in 0.2 ml collector tubes containing 20 μl lysis buffer (30 mM Tris-HCl pH = 8.0, 0.5% Tween-20, 0.5% NP-40 and 0.056 μAU Proteinase K) and kept on ice until digested at 56 °C for 60 minutes followed by 76 °C for 30 minutes, before being finally stored at −80 °C. For each animal, one or more pieces of ear skin were cut and processed as above to be used as germline controls.

##### Library preparation

Samples were processed using a recently developed enzymatic fragmentation-based library preparation method specifically designed to detect small amounts of DNA (low-input) without the need for whole genome amplification ^1^. Briefly, genomic DNA was purified by mixing 20 μl of the lysed samples with 50 μl Ampure XP beads (Beckman Coulter), and left for 5 minutes at room temperature. After magnetic separation, bead-DNA samples were washed twice with 75% ethanol and re-suspended in 26 μl TE buffer (10 mM Tris-HCl, 1 mM EDTA). For DNA fragmentation, samples were mixed with 7 μl 5x Ultra II FS buffer and 2 μl Ultra II FS enzyme (New England BioLabs), and incubated 12 minutes at 37 °C followed by 30 minutes at 65 °C. Samples were then incubated with a 30 μl ligation mix, 1 μl ligation enhancer (New England BioLabs), 0.9 μl nuclease-free water and 0.1 μl duplexed adapter (100 μM; 5⍰-ACACTCTTTCCCTACACGACGCTCTTCCGATC*T-3⍰, 5⍰-phos-GATCGGAAGAGCGGTTCAGCAGGAATGCCGAG-3⍰). Adapter-ligated libraries were then purified by mixing with 65 μl Ampure XP beads (Beckman Coulter) followed by magnetic separation and elution with 65 μl TE buffer. For the PCR reaction, 21.5 μl of the DNA samples were mixed with 25 μl KAPA HiFi HotStart ReadyMix (KAPA Biosystems), 1 μl PE1.0 primer (100 μM; 5⍰-AATGATACGGCGACCACCGAGATCTACACTCTTTCCCTACACGACGCTCTTCCGATC*T-31) and 2.5 μl iPCR-Tag (40 μM; 51-CAAGCAGAAGACGGCATACGAGATXGAGATCGGTCTCGGCATTCCTGCTGAACCGCTCTTCCGATC-3⍰), in which ‘X’ represents one of 96 unique 8-base indexes. PCR conditions were: 5 min at 98⍰°C, followed by 12 cycles of 30 s at 98⍰°C, 30 s at 65⍰°C and 1 min at 72⍰°C, and finally 5 min at 72⍰°C. Amplified libraries were purified by mixing with Ampure beads at a ratio 0.7:1, adjusted to 2.4nM with nuclease-free water.

##### DNA sequencing

A custom bait capture to target the exonic sequences of 192 genes frequently mutated in cancer was designed using Agilent SureDesign. The list of genes selected for ultra-deep targeted sequencing is shown below: *abcb11, abcc2, adam10, adam29, adcy10, aff3, ajuba, akap9, akt1, akt2, apob, arid1a, arid2, arid5b, asxl1, atm, atp2a2, atrx, aurka, b2m, bbs9, bcas3, bcl11b, bcr, braf, brca2, c1s, cacna1d, card11, casp8, ccnd1, cdc16, cdh1, cdkn2a, chuk, clgn, cnot1, cntnap4, cobll1, col12a1, copb2, cr2, crebbp, csmd2, ctcf, cul3, cyld, cyp2b13, dclk1, dclre1a, ddr2, dicer1, dmxl2, dnm2, dnmt3a, dst, egfr, eif2d, ep300, epha2, erbb2, erbb3, erbb4, ezh2, fat1, fat4, fbn2, fbxo21, fbxw7, fgfr1, fgfr2, fgfr3, flg2, flt3, fn1, gcn1l1, grin2a, grm3, gtf3c5, hist1h2bm, hmcn1, hras, huwe1, hydin, igsf1, insrr, iqgap1, irf6, kcnh5, kdm5b, kdm6a, kdr, keap1, kit, klrc3, kmt2c, kmt2d, kras, krt5, krtap4-9, loxhd1, lrp1, lrp1b, lrp2, ltf, maml1, mcm7, met, mrgprb4, mtor, myof, nf1, nf2, nfe2l2, nfkb1, nlrp12, notch1, notch2, notch3, notch4, nras, nsd1, nup214, opn3, pard3, pcdha5, pced1b, pde4dip, peg10, pign, pik3ca, pkhd1, plcb1, prex2, psme4, ptch1, pten, ptprt, rac1, rasa1, rb1, rbm46, rhbdf2, ripk2, ripk4, ros1, rpgrip1, rpl10, ryr2, sall1, scn10a, scn11a, scn1a, scn3a, setd2, setx, sgk3, sis, slc13a1, smad4, smarca4, smo, snx25, soat2, sox2, spen, st18, sufu, synm, taf2, tas2r102, tet2, tnr, trp53, trp63, trrap, tsc1, ttc27, usp24, usp26, usp9x, vhl, vmn2r81, vps13b, wnk1, zan, zfhx3, zfp39, zfp457, zfp521, zfp644* and *zfp750*.

Samples were multiplexed and sequenced on the HiSeq 2500 platform (Illumina) according to the manufacturer’s instructions, with the exception that we used iPCR-Tag (5⍰-AAGAGCGGTTCAGCAGGAATGCCGAGACCGATCTC-3⍰) to read the library index. Paired-end reads were aligned with BWA-MEM (v0.7.17, https://github.com/lh3/bwa) ^23^ with optical and PCR duplicates marked using Biobambam2 (v2.0.86, https://gitlab.com/german.tischler/biobambam2, https://www.sanger.ac.uk/science/tools/biobambam).

##### Mutation calling with deepSNV

In order to identify somatic mutations present in a small fraction of cells within the samples, we used the ShearwaterML algorithm from the deepSNV package (v1.21.3, https://github.com/gerstung-lab/deepSNV) to call for mutational events on ultra-deep targeted data ^2,3,24^. Instead of using a single-matched normal sample, deepSNV uses a collection of deeply-sequenced normal samples as the reference for variant calling that enables the identification of mutations at very low allele frequencies. For this study we used a total of 19 normal samples from 12 animals, providing a normal panel with a combined coverage of 7439x. A depth filter of 50x was used in the final variant call results.

##### Mutation burden

The average number of mutations per Mb in a given sample was estimated from the VAFs and the number of bases within the bait set with synonymous mutations (dubbed as synonymous footprint), as described before ^2,3^, and assuming no alterations in copy number as indicated by either the whole exome and genome sequencing data (**Extended Data Fig. 6**). Only the synonymous mutations were used for this calculation as the mutation burden can be inflated by the alterations in strongly selected genes when using targeted sequenced data.

##### Percentage of mutant epithelium

The size of mutant clones within each sample can be calculated by taking into account the area of the biopsy and the fraction of cells carrying a mutation within the sample, as described previously ^2,3^. We have determined that DEN treatment does not alter the copy number variation (**Extended Data Fig. 6**), therefore, the fraction of cells carrying a particular mutation can be estimated as twice the VAF of that mutation. The lower (=VAF) and upper (=2xVAF) bound estimates of the percentage of tissue covered by clones carrying non-synonymous mutations in a given gene was calculated for each sample ^2,3^. The fraction of epithelium covered by the mutant genes was then calculated from the mean of summed VAF (capped at 1.0) of all the samples.

### Mutational spectra and signature analysis

Mutational spectra for single base substitutions were plotted and compared to 65 known mutational signatures present in human cancers ^25^ using linear decomposition with the deconstructSigs R package (https://github.com/raerose01/deconstructSigs) ^26^. The mutational spectra across DEN-treated samples were highly consistent, precluding deconvolution into separate signatures (either known or de novo).

### Gene selection (dN/dS)

We used the maximum-likelihood implementation of the dNdScv algorithm (v0.0.1.0, https://github.com/im3sanger/dndscv) to identify genes under positive selection ^16^. dNdScv estimates the ratio of non-synonymous to synonymous mutations across genes, controlling for the sequence composition of the gene and the mutational signatures, using trinucleotide context-dependent substitution matrices to avoid common mutation biases affecting dN/dS. Values of dN/dS significantly higher than 1 indicate an excess of non-synonymous mutations in that particular gene and therefore imply positive selection, whereas dN/dS values significantly lower than 1 suggest negative selection. In our experimental set up, only mutations that reach a minimum size are detected by deepSNV. Therefore, a significant value of dN/dS>1 indicates that a clone that acquires a non-synonymous mutation in that particular gene will have a higher probability to reach a detectable size as compared to a synonymous mutation in the same gene. Hence, genes with dN/dS>1 are considered drivers of clonal expansion.

### Whole exome sequencing of surviving esophageal tumors

#### Sample preparation and imaging

Mice were treated with DEN in drinking water 3 times a week for 8 weeks as described above. 9-18 months after DEN removal mice were culled and the esophagus harvested. Tissues were incubated for 2–3 h in 5 mM EDTA at 37 °C before removing the submucosa from the epithelium as described above. The EEs were immuno-stained against KRT6 to detect the tumors and imaged on a fluorescent scope equipped with the appropriate filters. The projected area of the tumors was measured using Volocity 3D Image Analysis Software.

#### Tumor isolation and sequencing

Tumors were manually cut under a fluorescent micro-dissecting scope (Leica Microsystems) using ultra fine forceps and micro-scalpels. Individual tumors were collected in low binding DNA tubes and digested in 3 μl RLT buffer (Qiagen Cat# 1048449) for 30min at room temperature. Digested samples were diluted 1:10 in water, separated in triplicates, transferred to 96-well plates and incubated 15 min at room temperature with Agencourt AMPure XP magnetic beads (Beckman Coulter Cat# A63881) at a 1:1 ratio. Beads with bound DNA were separated with a magnet and washed 3 times with 70% ethanol. DNA was re-suspended in 10 μl elution buffer and transferred to a new plate. Whole genome DNA was amplified using 1 μl polymerase enzyme from the illustra GenomiPhi V2 DNA Amplification Kit (GE Healthcare Cat# 25-6600-32) and 9 μl of sample with the following conditions: 95 °C for 3 min, 4 °C for 5 min, 30 °C for 1.5 hours and 65 °C for 10min. DNA was then purified by mixing with beads at a 1:0.6 DNA/beads ratio followed by 3 washes with 70% ethanol and eluted with 30 μl of elution buffer (Qiagen Cat# 19086). Whole-exome sequencing was performed using the Mouse_Exome_Targets baitset from the Wellcome Sanger Institute pipeline. Captured material was sequenced on Illumina HiSeq 2500 sequencers using paired-end 75bp reads.

#### Mutation calling and sequencing analysis

Substitutions were called using the CaVEMan (Cancer Variants through Expectation Maximization) variant caller (version v1.11.2, http://cancerit.github.io/CaVEMan) ^27^. Insertions and deletions were called using cgpPindel (version v2.2.4, http://cancerit.github.io/cgpPindel) ^28^. Mutations were annotated using VAGrENT (v3.3.3, https://github.com/cancerit/VAGrENT) ^28^. Mutations occurring outside of the exome were removed by retaining only mutations annotated with the effects ‘missense’, ‘nonsense’, ‘ess_splice’, ‘frameshift' or ‘silent’. The mutational spectra and signature of single clones was analyzed as described above.

### Copy number analysis of whole exome sequenced tumors

The whole exome sequencing results were used to analyze the copy number alterations of surviving tumors. We used a modified version of QDNAseq (https://github.com/ccagc/QDNAseq/) to call somatic copy number alterations ^29^. QDNAseq was adapted to use only so-called “off-target” reads from exome sequencing, which yield an even, shallow coverage across the genome ^30^. To obtain off-target reads, we first counted all reads per 1Kb bins along the genome, then removed any bin that overlaps with the start/end coordinates of a protein-coding gene (Ensembl version GRCm38.p6) and mapped the remaining onto bins of 1Mb. Read counting was performed separately for each tumor sample and its matched control, after which the two coverage tracks were transformed into a log ratio track (commonly referred to as “logr”). The transformation occurs similarly to that of the Battenberg copy number caller ^31^: first the tumor bin-counts were divided by the control bin-counts to obtain the coverage ratio; the coverage ratio was then divided by the mean coverage ratio and finally the log2 was taken to obtain logr. The regular QDNAseq pipeline was then applied to adjust the logr for mappability and GC-content, and the genome segmented into regions of constant signal. Each segment was then subjected to a significance test to obtain copy number alterations. A t-test was applied between the logr of the segment and a logr of 0 (the expected logr if the segment is not altered), with the standard deviation of both distributions set to that of the observed data. A segment was not tested if its logr fell within [-0.05, 0.05] to conservatively account for remaining noise. The t-test had different power between segments, as these differed in size and therefore in the number of bins from which logr data was drawn. To account for this difference, we sampled up to 500 bins without replacement from the segment to perform the test. A one-sided t-test was then applied for gains and losses separately and p-values were adjusted for multiple testing using Bonferroni correction.

As every tumor was sequenced three times we applied a pipeline that used these triplicates to obtain confident copy number calls for each tumor. First, the method described above was applied to each sample and matched control, and we only kept alterations if they were called in all three samples that represent a tumor. This yielded 5 calls across 5 tumors. A tumor, however, had only a single matched control. This single control was used to normalize the data for all three triplicates of a tumor and could introduce false positive calls. We reasoned that a true alteration would remain when a sample is run against another control. We therefore randomly matched samples with a different control that was matched for sex, requiring each sample in a triplicate to be matched with a different control. The copy number pipeline was applied to each sample part of the 5 tumors that were found to contain an alteration in the first step, and again kept calls obtained in all three triplicates. This reduced the number of calls to 2, spanning 2 tumors.

### Whole genome sequencing of 1 year post-DEN esophageal neoplasms

#### Sample preparation and sequencing

Wild type mice were treated with DEN and the esophagus collected 1 year later. EE were stained with Dapi and KRT6 to label the tumors and imaged by confocal microscopy. Neoplasms were manually cut out under a fluorescent dissecting microscope. Samples were digested and DNA isolated as above (see WES methods section) and subjected to whole genome sequencing (median coverage 3.13x), using 150-base-pair clipped reads, on a HiSeq 4000 platform (Illumina).

#### Copy number analysis

We applied a modified version of QDNAseq (https://github.com/ccagc/QDNAseq/) that takes into account the control that is available for each sample for an additional normalization step, to obtain calls where the coverage log ratio is significantly different from a normal diploid state. Inspection of the results revealed a number of calls in a distinct region within chromosome 12 and on chromosome X in samples where the control was sample MD5921b. A run of control samples against each other also revealed frequent calls on chromosome 12, spanning a region 5Mb (bp 18,200,001 to 23,300,000), which was subsequently blacklisted. We further used the largest call (1.15Mb) in this run of control samples to establish a threshold on the minimum genomic region to be covered, and kept calls only if they exceed it (3 calls filtered). Inspection of control sample MD5921b revealed unusually low coverage, which suggests the gains called in samples where this control was used during analysis are likely false positives. We therefore removed calls on chromosome X in these samples (14 calls filtered). After filtering, the final set of alterations contains a gain that covers the whole of chromosome 19 in sample MD5924e and a focal loss on chromosome 14 in sample MD5928e (**Extended Data Fig. 6a**).

### Detection of apoptosis by activated caspase-3 staining

Wild-type mice received DEN in drinking water for eight weeks as described above, and culled 10 days after the treatment stopped. Esophagus were collected, fixed, and immuno-stained to label active caspase-3 and KRT6. Tissues were imaged by confocal microscopy using a x40 objective and the presence of caspase-3 positive cells within the tumors or in the surrounding normal epithelium was analyzed with Image J.

### EdU lineage tracing assay

EdU (5-ethynyl-2⍰-deoxyuridine) incorporates into dividing cells and therefore quantification of EdU positive cells can be used as a direct measure of cell proliferation in the tissue. Mice received DEN in drinking water for 8 weeks as described above. 10 days after finishing DEN treatment mice were administered 10 μg of EdU (i.p.) and the esophagus collected 1h later. Tissues were peeled, fixed and the incorporated EdU detected using a Click-iT EdU imaging kit (Life technologies Cat# C10086) according to the manufacturer’s instructions. Tissues were imaged by confocal microscopy using a x40 objective and the presence of EdU positive cells in the tumors and their adjacent normal epithelium analyzed using IMARIS software.

### In vivo transgenic label-retaining cell assay

*Rosa26^M2rtTA^/TetO-HGFP* mice were used to measure the rate of cell division in DEN generated tumors as compared to their adjacent normal epithelium. These animals are double-transgenic for a reverse tetracycline-controlled transactivator (rtTA-M2) targeted to the Rosa 26 locus and a *HIST1H2BJ/EGFP* fusion protein (Histone-Green Fluorescent Protein, HGFP) expressed from a tetracycline promoter element ^32^. Administration of doxycycline (Doxy, Sigma Aldrich Cat# D9891) induces the transient expression of HGFP, resulting in nuclear fluorescent labelling throughout the entire epithelium. When Doxy is withdrawn, HGFP is no longer expressed and is diluted lineally by half after every cell division cycle (**Fig. 3b**). As a result, the decline in fluorescence intensity can be used to calculate the cell division rate. *Rosa26^M2rtTA^/TetO-HGFP* mice received DEN for eight weeks as described above. During the last 4 weeks of DEN treatment, Doxy (2mg/ml) was also supplied in the drinking water. After DEN-Doxy treatment, mice were culled and esophagus collected at times t= 0 (immediately after the treatments stopped), t = 10 days or t= 30 days. Tissues were peeled, fixed and stained (nuclei and KRT6) as detailed above, and imaged on a confocal microscope using a 40x objective. The intensity of HGFP in the tumors and the adjacent normal epithelia was analyzed using ImageJ.

### Leukocyte density in tumors and normal epithelium

Wild-type mice were treated with DEN for 2 months and the esophagus collected 10- or 30-days after treatment. The esophageal epitheliums were processed and immuno-stained with anti-CD45 and anti-KRT6 antibodies to visualize leukocytes and tumors, respectively, and imaged on a confocal microscope. Typical settings for acquisition of multiple z stacks were 2.5μm z-step size, zoom x1, optimal pinhole, line average 4, scan speed 400⍰Hz and a resolution of 1,024 × 1,024 pixels using a 10X HC PL Apo CS Dry objective with a 0.4NA. Visualization and image analysis were performed using IMARIS (bitplane) and ImageJ software. Images, with the tumors placed at the center of a 300 x 300 microns field of view, were used to quantify the number of leukocytes within and outside the tumors. This number was normalized by the area of the tumor or the surrounding normal tissue (calculated by subtracting the area of the tumors to the total area).

### Permutation analysis of leukocyte location within tumors and normal tissue

To investigate whether the number of immune cells was enriched or excluded within the tumors, or instead followed a random distribution, we use a permutation analysis based on the experimental measurements of leukocyte density in tumors and normal epithelium obtained as detailed above. For this, tumors were assigned a circular shape, with an area matching that measured experimentally. Immune cells (CD45^+^ cells) were also assumed to be circular with a diameter of 8 microns. For each field of view, the location of the immune cells was left intact while the location of the tumor was randomly shuffled (ensuring the entirety of the tumor remained within the boundaries of the field of view), and the number of immune cells in contact with the tumor was counted (Extended Data Fig. 7h). This was repeated 1000 times to produce the expected distribution assuming no association between tumor and immune cell locations.

### Induction of DN-Maml1 clones

*Ahcre^ERT^R26^flDNM−GFP/wt^* mice were used for induction of the dominant negative mutant of Maml1 (DN-Maml1) ^9^. This mutant inhibits Notch intracellular domain induced transcription, therefore disrupting the Notch signaling pathway. It is also fused to GFP, which allows for clonal labelling of the mutant. Expression of DN-Maml1 can be genetically induced following treatment with ß-napthoflavone (BNF, MP Biomedicals Cat# 156738) and tamoxifen (TAM, Sigma Aldrich Cat# N3633). Specifically, transcription of the Cre mutant estrogen receptor fusion protein (CreERT) is induced following intraperitoneal (i.p) BNF injection. A subsequent i.p injection of TAM is necessary in order for the CreERT protein to gain access to the nucleus and excise the loxP flanked “STOP” cassette resulting in the expression of DN-Maml1 tagged to GFP. As the switch occurs at the gene level, the descendants of the originally labelled cell (clones) will also constitutively express DN-Maml1, and can be visualized by fluorescent microscopy.

Mice received DEN for 8 weeks as described above. 10 days after DEN withdrawal, mice were given a single injection of BNF (80⍰mg⍰kg) and TAM (1⍰mg), or left un-induced. Esophagus were collected 30 days post-DEN treatment (20 days after induction of DN-Maml1 clones). Whole tissues were processed, stained with anti-GFP and KRT6 antibodies and imaged on a confocal microscope as described above. The percentage of esophageal epithelium occupied by DN-Maml1 mutant clones and the number of tumors within and outside DN-Maml1^+^ areas were measured using Volocity 3D Image Analysis Software (Perkin Elmer) or IMARIS software.

### Percentage of tumors eliminated by DN-Maml1 clones

The percentage of tumors removed by expanding DN-Maml1 clones was calculated as follow: % of tumors eliminated = (1-T/(DNneg)) 100. Where *T* is the total number of tumors per mm^2^ of esophageal epithelium and is the number of tumors per mm^2^ of DN-Maml1 negative epithelium within the same tissue.

### In vivo treatment with the *γ*-secretase inhibitor DBZ

Wild-type mice received DEN in drinking water for 8 weeks as described above. 10 days after DEN withdrawal, mice were injected i.p. with the γ-secretase inhibitor DBZ (S2711; Selleckchem) at 30μmol/Kg of bodyweight, using a stock solution of 2.8mg/ml in 0.5% hydroxypropyl methylcellulose (Methocel 65HG, 64670-100G-F, Sigma) and 0.1% Tween-80 in water. Control animals were injected with vehicle solution. Mice received DBZ or vehicle every three days, a total of 5 injections, and were culled 2 weeks after initiating the treatment (24 days post-DEN). Esophagus were collected and epitheliums from control and DBZ-treated mice processed and stained with KRT6 antibodies. Whole-mounted esophageal epitheliums were imaged by confocal microscopy and the number of tumors per mm^2^ of epithelium measured using IMARIS software.

### Fitting of model to data

See **Supplementary information**. The parameter sets which best fit the data were found using Approximate Bayesian Computation based on sequential Monte Carlo sampling (ABC-SMC) ^33^ implemented in the python package pyABC ^34^. This method randomly generates parameter combinations, then uses summary statistics to compare the model’s output using those parameters to the observed data, and rejects parameter combinations for which the model’s output is too dissimilar to the data. Multiple generations are run with increasingly stringent thresholds for acceptance, with each new generation of parameters based on the accepted parameter combinations from the previous generation. In this manner, the method identifies the regions of parameter space that produce model outputs most similar to the observed data. We ran 25 generations with a population of 10000 (the number of accepted parameter combinations in each generation). For the summary statistic, we used the total squared distance of the mean model prediction from the mean observed tumor density. For the tumor drift model with no interaction with mutant clones in surrounding normal tissue, the fit was to the tumor density following DEN-treatment. For the model including clone interaction, the fit was to the tumor density following DEN-treatment and the tumor density in the DBZ experiment. Uniform distributions using the bounds in **Supplementary information Table A** (see **Supplementary information**) were used as the initial prior distributions of the parameters.

### Statistical analysis

Data are expressed as mean values ± s.e.m. unless otherwise indicated. No statistical method was used to predetermine sample size. The experiments were not randomized. The investigators were not blinded to allocation during experiments and outcome assessment.

## Data availability

Individual data sets are available in Supplementary Tables 1–8. The accession number for the DNA targeted sequencing of 10 days and 1 year tumors is ENA: ERP022921. Accession numbers for WES and WGS of tumors are ENA: ERP015469 and ENA: ERP122780, respectively.

## Code availability

Code is available at https://github.com/michaelhall28/Colom_lesions.

## Acknowledgments

We thank Esther Choolun, Tom Metcalf and staff at the MRC ARES and Sanger RSF animal facilities for technical support. This work was supported by grants from the Wellcome Trust to the Wellcome Sanger Institute (098051 and 296194) and Cancer Research UK Programme Grants to P.H.J. (C609/A17257 and C609/A27326). B.A.H. and M.W.J.H. are supported by the Medical Research Council (Grant-in-Aid to the MRC Cancer unit grant number MC_UU_12022/9 and NIRG to B.A.H. grant number MR/S000216/1). A.H. benefited from the award of an EMBO long term fellowship. M.W.J.H. acknowledges support from the Harrison Watson Fund at Clare College, Cambridge. B.A.H. acknowledges support from the Royal Society (grant no. UF130039). S.D. benefited from the award of an ESPOD fellowship, 2018-21, from the Wellcome Sanger Institute and the European Bioinformatics Institute EMBL-EBI. K.T.M. benefited from the support of the Chan Zuckerberg Initiative. We are grateful to the Cambridge Biorepository for Translational Medicine for access to human tissue.

## Author contributions

B.C., M.P.A. and A.H. designed experiments. B.C., A.H., M.P.A., G.P. and D.F.A performed experiments and analyzed data. B.C., M.W.J.H., S.C.D., C.K., R.K.S. and S.H.O. analyzed sequencing data. M.W.J.H. developed the mathematical modelling. K.T.M., K.S.P., and J.C.F. collected human samples. B.C., M.W.J.H., and P.H.J. wrote the paper. B.A.H., M.G. and P.H.J. supervised the research.

**Supplementary information is available for this paper:** see below

**Correspondence and requests for materials should be addressed to P.J.H.**

## Supplementary information

This document describes the whole exome sequencing (**section 1**) and analysis of chromosomal number alteration (**section 2**) of surviving tumors, as well as the experimental work to explore potential mechanism of elimination of early tumors such as tumor cell apoptosis and loss of proliferation (**section 3**), and the immune system (**section 4**). The theory and mathematical modeling of tumor dynamics is then set out (section 5).

### Section 1. Whole exome sequencing (WES of surviving tumors

The dN/dS analysis of targeted sequenced samples indicated that 5 out of the 192 genes analyzed, *Atp2a2, Notch1, Notch2, Chuk* and *Adam10* are significantly (q<0.05) positively selected in tumors persisting 1 year after DEN treatment (**Fig. 2c**). To explore whether other genes not included in our targeted sequencing bait set were selected in these samples, we performed whole exome sequencing (WES) of persisting tumors.

Wild type mice were treated with DEN and esophagus collected 9-18 months later (**Extended Data Fig. 5a**). EE were stained with Dapi and antibodies against KRT6 to label neoplasms and imaged by confocal microscopy. 49 tumors from 16 tissues were identified, manually cut out under a fluorescent dissecting microscope and digested. Genomic DNA was isolated from each sample and split into three pools, each of which underwent independent whole genome amplification (WGA) and WES to an average cumulative coverage of 1130x (**Extended Data Fig. 5a-b**). To exclude artefactual SNVs generated during WGA, only mutations shared by all three amplified triplicates were included in the subsequent analysis.

After applying these criteria a total of 32,736 mutations were identified, including silent, missense, nonsense and splice mutations and indels (Supplementary Table 6). The median number of SNVs/exome per isolated tumor was 669, ranging from 56 to 1256, with an average variance allele fraction (VAF) of 0.233 ± 0.124 (mean ± SD) (Extended Data Fig. 5c-d). The spectrum and functional impact of mutations was consistent with the results obtained with the targeted sequencing approach (Extended Data Fig. 5e-f). Finally, dN/dS analysis indicated that *Atp2a2* and *Notch1* were positively selected in these samples (Extended Data Fig. 5g and Supplementary Table 7).

### Section 2. Analysis of DNA copy number alterations (CNAs in 1 year tumors

CNAs are rare in normal human esophageal epithelium but common in esophageal cancers ^2,5,35^. Thus, we explored whether the presence of CNAs would be a feature of the surviving tumors at 1 year post-DEN treatment.

We initially performed whole genome sequencing (WGS) of 64 tumors from 9 esophagus, with a median coverage of 3.13x. Control samples obtained from the same animals were used for an additional normalization step. After applying some custom filters (**see Methods**) only two tumors showed evidence of chromosomal alterations, where the coverage log ratio was different from a normal diploid state. These include a gain of the whole chromosome 19 and a focal loss on chromosome 14 (**Extended Data Fig. 7a**).

We next interrogated our WES dataset to look for CNAs, to ensure the above WGS was not underpowered due to the polyclonality of these samples ^15^ (**Section 1**). Somatic CNAs were called from the WES tumors as reported before ^11^. In line with the results obtained with WGS, the analysis of WES samples showed negligible presence of CNAs in the surviving tumors. Only 2 of the 49 tumors analyzed exhibited evidence of alterations, particularly in Chromosomes 7 and 16 (**Extended Data Fig. 7b-c**).

### Section 3. Early neoplasms are not eliminated by increased apoptosis or decreased proliferation of tumor cells

Here we set out how we investigated apoptosis and the rate of proliferation in murine normal and neoplastic EE.

To determine whether the elimination of early tumors was related to increased apoptosis of tumor cells, wild-type mice were treated with DEN for two months and the esophagus collected 10 days after the last treatment. Tissues were then immuno-stained against KRT6, to label tumors, and activated Caspase-3 as a maker of apoptosis, and imaged by confocal microscopy (**Fig. 3a**) In line with our previous observations we found negligible activated Caspase-3 positive cells in the DEN-treated normal epithelium (~0.04% of cells) ^11^. Importantly, no Caspase-3 positive cells were detected within the tumors, suggesting that early neoplasms are not lost through apoptosis.

Another potential mechanism for the elimination of early tumors could be the loss of proliferating tumor cells. To assess this, we first injected EdU (5-ethynyl-2⍰-deoxyuridine) into mice, 1 hour before tissue collection (**Extended Data Fig. 7a**). EdU incorporates into replicating DNA and labels cells in S phase of the cell cycle. We found that cells within the tumors incorporate EdU to a similar extent than their surrounding normal epithelium (**Extended Data Fig. 7b**). We next measured the rate of cell division in transgenic assay using *Rosa26^M2rtTA^/TetO-HGFP* mice. In this strain, expression of a stable Histone-Green Fluorescent Protein (HGFP) is regulated by a Doxycycline-responsive transgenic promoter. When Doxycycline is withdrawn, levels of the stable HGFP protein fall at a rate dependent on the rate of cell division **Fig. 3b**) ^13,15,32^. *Rosa26^M2rtTA^/TetO-HGFP* mice received DEN for 4 weeks followed by DEN + Doxycycline for another 4 weeks. Animals were culled and tissues collected at times 0, 10 days and 30 days after DEN + Doxycycline withdrawal (**Fig. 3c**). Analysis of the intensity of HGFP in tumors gradually decreased with time (**Fig. 3d and Extended Data Fig. 7c**). Importantly, the rate of HGFP loss at each time point was not significantly different between the tumors and the adjacent normal epithelium, indicating that cells within the tumors divided at a similar rate than the surrounding normal epithelium.

From these results we conclude that micro-tumors were not lost due to apoptosis or abnormal proliferation of tumor cells.

### Section 4. Early neoplasms are not eliminated by the immune system

We next explored whether micro-tumors were recognized and eliminated by the immune system. If this was the case, then tumors would not be removed as efficiently in mice with a defective immune system. We tested this using *NOD.Cg-Prkdc^scid^ Il2rg^tm1Wjl^/SzJ* ^22^ mice, a strain that lacks typical anti-tumor immune responses due to a complete absence of B and T lymphocytes and NK cells, as well as defective dendritic cells and macrophages. *NOD.Cg-Prkdc^scid^ Il2rg^tm1Wjl^/SzJ* and wild-type control mice received DEN for 8 weeks, and tissues were collected 10 and 30 days post-DEN withdrawal, stained and analyzed by confocal microscopy (**Fig. 3e**). There was no significant difference between strains in the number of tumors lost (**Fig. 3f**). This indicates that tumors are removed even if the immune response is severely compromised. Of note, the lack of a significant difference in the number of tumors at 10 days also suggests that the immune system does not have a critical role in the formation of tumors in this model.

To further explore the relationship between immune cells and early tumors we analyzed the spatial localization of leukocytes (CD45^+^ cells) in EEs from DEN-treated wild-type mice collected 10 days and 30 days post-DEN withdrawal (**Extended Data Fig. 7d**). The results show a positive correlation between the size of the tumors and the number of immune cells within them (**Extended Data Fig. 7e**). Furthermore, our analysis indicates that the number of immune cells normalized by tumor area positively correlates with that in the adjacent normal tissue (**Extended Data Fig. 7f**). Consequently, no significant differences were found in the average number of immune cells present within the tumors and at the surrounding normal EE at either the 10 or 30 day time points (**Extended Data Fig. 7g**). These results argue that the presence of immune cells within early neoplasms follows the same pattern than the adjacent normal tissue, suggesting that immune cells are not able to detect the tumors at this early stage.

Finally, we used a different approach to investigate whether immune cells were enriched, excluded or behaved neutrally with respect to the early tumors. For this purpose, we performed a permutation analysis taking advantage of the known spatial localization of the immune cells in the images from the above experiments. For each field of view, the coordinates of the immune cells were fixed, the tumor randomly shuffled within the image and the number of immune cells in contact with the “phantom” tumors quantified (**Extended Data Fig. 7h**). The observed number of immune cells in contact with the tumors differed slightly from the calculated numbers due to the assumed circularity of the tumors and immune cells (**Extended Data Fig. 7i** and Methods). Despite this, the number of immune cells in contact with tumors was not significantly different from a random distribution (**Extended Data Fig. 7j**). We conclude that immune cells are not enriched or selected but behave neutrally with respect to early micro-tumors.

All together, these results argue that immune cells are not responsible for the early elimination of tumors.

### Section 5. Theory and mathematical modeling

#### Modelling aims

Here we describe a general model of tumor loss that allows us to make testable predictions without requiring details of cell dynamics in tumors or mutant clones in the epithelium. We are interested in particular in whether the tumors are lost due to tumor-intrinsic mechanisms, or if the tumors are removed by mutant clones in the adjacent normal epithelium. As we discuss the factors affecting tumor loss, the principles of the model are described in mathematical terms so that we can later construct a numerical version of the model. This numerical version of the model is intended as a demonstration of the model principles, rather than a parameter- or model-fitting exercise.

In **section 5.1**, we briefly summarize previous results regarding growth and competition of mutant clones in normal EE. In **section 5.2**, we describe a previously proposed stochastic model of tumor dynamics and apply it to the tumor density following DEN-treatment. In **section 5.3**, we show that tumor loss is affected by the induction of highly fit mutant clones in the surrounding tissue. In **section 5.4**, we show how reducing the competitive imbalance between tumors and highly fit mutant clones in the normal tissue increases the number of surviving tumors. In **section 5.5**, we discuss how certain tumor genotypes may have higher survival chances due to a resistance to displacement by mutant clones in the normal tissue. In **section 5.6**, we substitute simple mathematical equations into the model to numerically illustrate the principles described in the previous sections. **Section 5.7** is a brief summary of our conclusions

##### 5.1 Clonal competition in esophageal epithelium

Before examining the behavior of tumors in the tissue, we first summarize previously published results relating to competition of mutant clones in phenotypically normal murine EE ^11^.

The EE is maintained by proliferating cells in the basal layer (**Extended Data Fig. 1a**). These cells divide stochastically (randomly) (**Extended Data Fig. 1b**), meaning that, if we track the offspring of basal cells, some cell lineages will grow into multicellular clones of the original cell, and other lineages will be lost as all basal cells differentiate ^13^. A mutation in a basal cell may convey a growth advantage, biasing cell fate towards producing more dividing daughter cells and promoting the growth of a mutant clone ^9–12,36^ (**Extended Data Fig. 1c**).

When the tissue contains multiple mutant clones, the clones compete for the limited space in the tissue ^11^ (**Extended Data Fig. 1d**). Fitter clones are able to displace less fit clones, but once a clone is surrounded by clones of similar competitive fitness it returns towards more neutral growth ^11^ (**Extended Data Fig. 1d**). The expansion and survival of a mutant clone therefore depends not only on its own fitness, but the fitness of its neighbors ^11^.

Following DEN treatment, the density of tumors in the EE falls rapidly (**Fig. 1d**). Tumors are not lost due to chromosomal copy number alterations (**Extended Data Fig. 7** and **section 2** above), apoptosis (**Fig. 3a** and **section 3** above), loss of proliferation (**Fig. 3d, Extended Data Fig. 7b-c** and **section 3** above) or interaction with the immune system (**Fig. 3f, Extended Data Fig. 7d-j** and **section 4** above). The tumors grow amongst a dense patchwork of highly fit clones competing for their place in the tissue. In the following sections we explore whether the tumors survival is affected by the mutant clones around them.

##### 5.2 Stochastic model of tumor dynamics

In a previous study we found that the growth of high grade dysplasias (HGDs) in the esophagus of DEN and Sorafenib treated mice was consistent with a stochastic model of cell dynamics ^15^. In this model, proliferating cells divide to form a pair of proliferating daughter cells, a pair of non-dividing daughter cells, or one cell of each type (**Extended Data Fig. 1b-c**). In the normal tissue, the probabilities of each symmetric division type are balanced, so that the total number of proliferating cells remains approximately constant (**Extended Data Fig. 1b**. In the HGDs, there was a bias towards producing more dividing than differentiated cells, causing the average size of tumors to increase over time (**Extended Data Fig. 1c**) ^15^.

Due to the stochastic nature of the process, some tumors expand in size, while in others all basal cells differentiate and the tumor is shed and lost from the tissue ^15,37^. This variation in outcome due to random chance is known as drift. If we assume that the fully differentiated tumors are lost from the tissue, then the survival probability of a tumor is given by the equation ^38^:

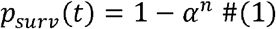

where *n* is the initial number of proliferating cells in each tumor (assumed here to be the same for all tumors), and

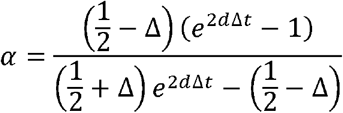

Where *d* is the symmetric division rate of tumor cells, Δ is the probability imbalance between symmetric division and symmetric differentiation (**Extended Data Fig. 1c**), and t is time. Fitting equation 1 to the tumor density following DEN-treatment results in a steep initial drop in tumor numbers followed by a slower downward trend, consistent with the experiment (**Supplementary information Figure A**). The median accepted parameters found upper bound) *d*=0.33/day (0.27, 0.36), Δ=0.003 (0.001, 0.005), *n*=1.2 (1.0, 1.5), and initial from fitting the model to the data (**see Methods**) were (95% credible interval lower bound, tumor density (immediately following DEN-treatment)=4.8/mm^2^ (4.1, 5.6), though these parameters should not be interpreted as estimates of the true biological values (**see sections below**).

**Supplementary information Figure A.**
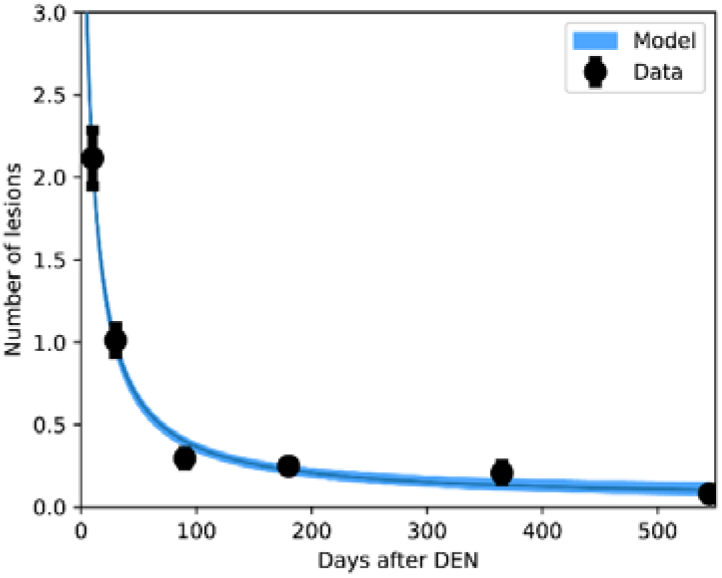
Tumor drift model (with no interaction with mutant clones in surrounding normal tissue) fit to tumor density (# tumors per mm^2^ of EE) following DEN-treatment (see **Methods**). The mean value and range between the minimum and maximum values of the models run with the accepted parameters are shown.

However, we will see in the sections below that this model is not capable of explaining the results seen in the full range of experiments, and therefore must be rejected (or at least adjusted) to account for the clones in the surrounding tissue.

##### 5.3 Elimination of tumors by highly competitive mutant clones in the surrounding normal epithelium

Although the stochastic model of tumor drift defined above is sufficient to describe the pattern of tumor loss following DEN-treatment, it does not rule out alternative causes of tumor loss. The normal EE surrounding the tumors contains a patchwork of competing mutant clones (see **section 5.1**). Therefore, we speculated that, like clones in the surrounding normal tissue, tumors may be displaced by highly fit mutant clones.

We considered a general model in which a highly competitive mutation, *M*, is induced in the tissue following DEN treatment. We assume that clones of the mutant *M* are able to remove tumors that they encounter (**Extended Data Figs. 8a and 9c**). Let *p_M_(t)* be the survival *p_other_* be the survival probability of a tumor based on all other sources of tumor loss (such probability of a tumor assuming that the mutant M is the only cause of tumor loss. Let as drift – see **section 5.2**, but may include tumors removed by DEN-created clones – see **section 5.4** below). For simplicity, we assume that tumor loss due to the mutant M is independent of all other causes of tumor loss. The combined survival probability of a tumor is then given by

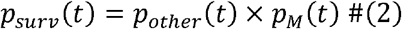

Let *M(t)* be the proportion of tissue covered by the mutant at time *t*. We make the following additional assumptions for the sake of simplicity:

1. The proportion of tissue covered by *M* increases monotonically.
2. Tumors are spread randomly across the tissue.
3. Tumors are removed instantly with probability 1 when the mutant *M* colonizes the location of the tumor.

This means

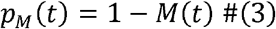

and the combined probability of tumor survival is then given by

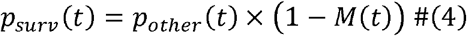

To make it easier to compare mice in which the initial density of tumors may vary, we looked at the proportion of lesions eliminated (*PLE*) by *M*, using the tumor density in the non-M-mutant regions of the tissue to estimate the tumor density in the full tissue in the absence of *M*.

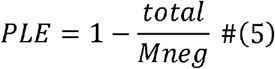

where *total* is the density of tumors over the full tissue, and *Mneg* is the density of lesions in the M-negative region.

The expected PLE for two models - where the *M* mutant removes lesions it encounters (*PLE_M_*), and where tumor loss is independent of the mutant in the surrounding tissue (*PLE_¬M_*), are

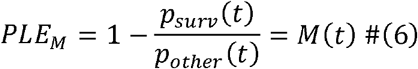

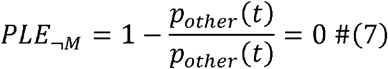

In other words, if *M* clones are able to remove tumors in a similar manner to which highly fit clones are able to displace weaker clones in the normal tissue, then we will see a reduction in tumor numbers proportional to the spread of the *M* mutant. By definition, models in which tumor survival is independent of surrounding tissue (*PLE_¬M_*), such as the stochastic tumor drift model described in **section 5.2**, predict that tumor numbers will be unaffected by the spread of the *M* clones (**Supplementary information Figure B**).

**Supplementary information Figure B.**
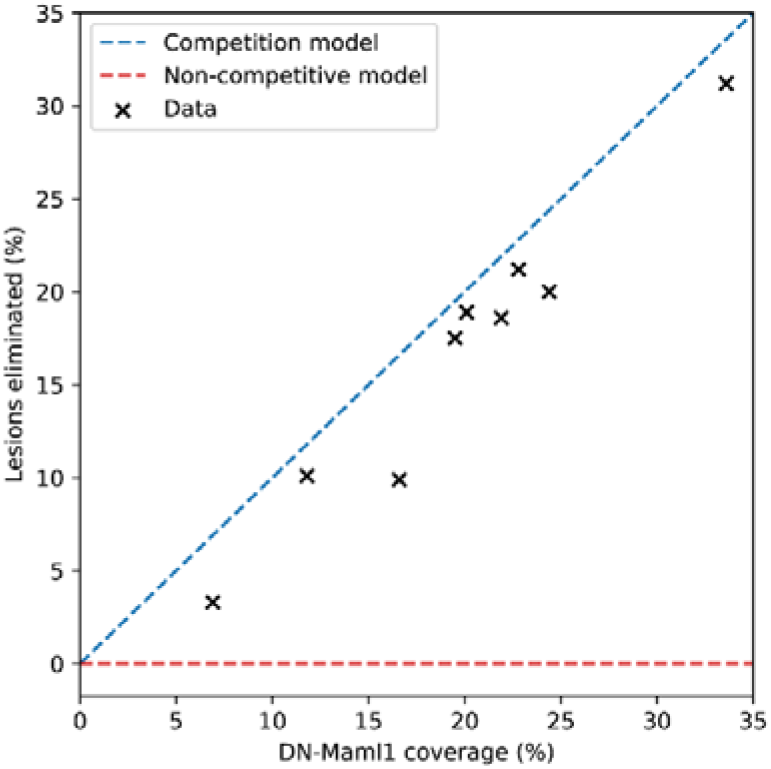
Tumor loss when highly fit mutant clones (such as DN-Maml1 clones) are induced in EE following DEN-treatment. Predictions of a model in which the induced mutant clones immediately remove lesions they encounter (blue) and a model in which tumor loss is independent of clones in the surrounding tissue (red).

To test these predictions, we used an inducible DN-Maml1 mutation that prevents Notch signaling and forms rapidly expanding clones when induced in murine EE ^11^. The data shows a strong correlation between DN-Maml1 clone spread and tumor loss, indicating that DN-Maml1 clones are indeed removing tumors from the tissue (**Fig. 4c-d, Supplementary information Figure B**).

In the experiment, there remained a small number of tumors in close contact with DN-Maml1 mutant regions (**Fig. 4e**). This may indicate that there is a lag time between contact with DN-Maml1 clones and tumor removal and that the surviving tumors seen in the DN-Maml1 areas are in the process in being removed (**Fig. 4e and Extended Data Fig. 9a-c**). It may also be the case that a small proportion of tumors are able to survive despite the competition with DN-Maml1 clones (see section 5.5 below).

##### 5.4. Reducing competitive imbalance

Now that we have shown that the induction of a highly fit mutant following DEN treatment can eliminate tumors, we asked whether mutant clones already present in the DEN-treated tissue are able to remove tumors too. Fit clones present in the normal epithelium might be able to out-compete the tumors, eliminating them from the tissue. By removing the competitive advantage of those clones, we can examine the impact they are having on tumor survival.

We assume there is a type of highly fit mutant clone, *N*, in the DEN-treated tissue that is able to remove tumors. As in the section above (Equation 2), we assume that the removal of clones by mutant clones *N* is independent of other causes of tumor loss. Tumor survival probability is given by

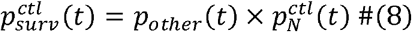

where, similar to above, *p_other_* (*t*) is the survival probability of a tumor based on all sources of tumor loss other than elimination by *N* clones, and 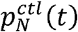 is the survival probability of a tumor assuming that the mutant *N* is the only cause of tumor loss. We assume that, without intervention, *N* clones spread progressively throughout the tissue and outcompete tumors they encounter, so 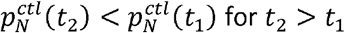.

If we can raise the fitness of the surrounding tissue and tumors to a similar level as *N* clones, this would both prevent the spread of the *N* clones across the tissue (reducing the number of tumors directly competing with *N* clones) and reduce the elimination of tumors that are already adjacent to *N* clones, as they will now be competing neutrally (Extended Data Fig. 10a). Assuming that we have an intervention that completely levels the fitness of *N* clones with the rest of the tissue and tumors, the loss of tumors due to *N* clones will cease during that period, i.e. if the intervention starts at time *t*_1_ and lasts until *t*_2_, 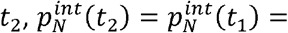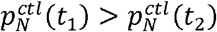 and

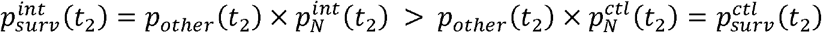

where, for the experiment in which the intervention is applied, 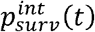 is the overall tumor survival probability and 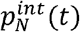 is the tumor survival probability related to the mutant N. Therefore, if we can remove or reduce the competitive imbalance between a mutant that can remove tumors and the rest of the tissue (including the tumors), then we should see an increase in surviving tumors compared to control experiments (**Extended Data Fig. 10b**).

Notch1 mutant clones dominate competition in the normal EE ^11^ and therefore is a good candidate mutation to test this prediction. The Notch inhibitor Dibenzazepine (DBZ) prevents Notch signaling and would affect all cells in the tissue, effectively raising the fitness of all Notch wild type clones and tumors to the level of the Notch mutant clones ^9^. The competitive advantage of Notch mutants is thus removed during the DBZ intervention. Furthermore, by raising all cells to a high background level of fitness, this may also reduce the relative fitness advantage conveyed by mutations which work independently of the Notch pathway ^39^.

We indeed saw the predicted increase in surviving tumors when DBZ is administered between 10 days (*t*_1_) and 24 days (*t*_2_) after DEN treatment (**Extended Data Fig. 10c-d and Supplementary information Figure C**). This suggests that competition from mutant clones in the surrounding tissue is removing a substantial proportion of the tumors lost in the first few weeks following DEN treatment.

**Supplementary information Figure C.**
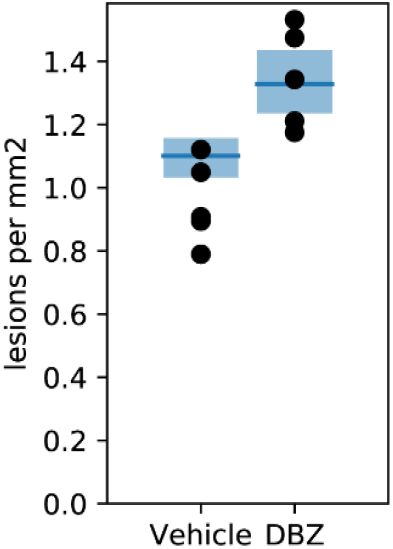
Numerical example of the model (see **section 5.6**) showing increased tumor survival by administration of DBZ as compared to vehicle control. Model fit to lesion density data using Approximate Bayesian Computation (see **section 5.6** and **Methods**). The mean value and range between the minimum and maximum values of the model run with the accepted parameters are shown.

##### 5.5 Selection pressure on tumors from competition with surrounding mutant clones

There are genetic differences between the tumors sequenced 10 days and 1 year after DEN treatment, as shown by the dN/dS ratios (**Fig. 2c**) and the proportion of tumors mutant for each selected gene (**Extended Data Fig. 4g**). This genetic change over time could be consistent with ongoing selection of mutant subclones within tumors ^40^. However, the elimination of early neoplasms by mutant clones in the surrounding tissue could also act as a selective pressure on tumors.

As described in **section 5.4** above, the competitive fitness of a tumor compared to the surrounding clones will affect the survival prospects of the tumor (**Fig. 4a**). If certain tumor genotypes have a competitive fitness comparable to or higher than the fittest mutant clones in the surrounding tissue, they would be more likely to survive. For example, *Notch1* is the dominant mutant gene in normal tissue, occupying almost the entire tissue 12 months after DEN treatment ^11^. We might expect that, similar to the *Notch1* mutant clones in the normal tissue, tumors which are also Notch1 mutant would be able resist displacement by the mutant clones in the surrounding tissue. Consistent with this, we saw an increase in the proportion of Notch1 mutant tumors from the 10-day to the 1-year time point (**Extended Data Fig. 4g**). We also see a large increase in the proportion of *Atp2a2* mutant tumors (**Extended Data Fig. 4g**), suggesting that these too may be able to resist displacement by clones in the surrounding tissue.

To explore this hypothesis, we expanded the model to include two tumor phenotypes: sensitive and resistant. Sensitive tumors can be removed by clones in the surrounding tissue, and resistant clones cannot. The probability of tumor survival is then given by

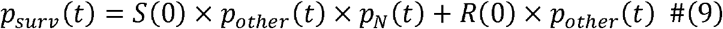

where *S*(0) and *R*(0) are the proportions of tumors at day 0 following DEN treatment which are sensitive and resistant respectively.

The proportion of surviving tumors which are resistant to displacement by mutant clones, *R*(t), is given by

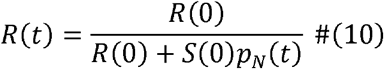

Assuming that the number of sensitive and resistant tumors are non-zero, and that, as we assumed earlier, *p_N_(t)* is a decreasing function of time, then the proportion of surviving tumors which are resistant will increase over time.

The increasing proportion of the surviving tumors which are *Notch1* and/or *Atp2a2* mutant is consistent with the hypothesis that these mutations may increase the tumor’s ability to resist displacement by clones in the surrounding tissue. The presence of a subset of tumors which are able to resist displacement by clones also could explain why a small fraction of tumors are able to survive over 1 year after DEN treatment (**Fig. 1d**) when the surrounding normal tissue is almost entirely populated by highly fit mutant clones ^11^.

##### 5.6 Numerical model example

So far, we have mostly defined properties of the tumor survival probabilities rather than given specific equations for their values. This has allowed us to make testable predictions without requiring the details of the clonal or tumor dynamics to be defined.

Here we substitute feasible functions into Equation 9 to construct a numerical expression of the model. The functions are intended to be simple and introduce only a small number of model parameters, and the purpose here is to illustrate the concepts described in the previous sections rather than accurately and verifiably model the data.

Firstly, we need to define *p_other_*, the non-clone related probability of tumor survival. As we have shown that apoptosis and abnormal proliferation of tumor cells, and the immune above), we assume that *p_other_* is simply the survival probability based on drift (Equation 1). system are not contributing to tumor loss (**Fig. 3, Extended Data Fig. 7 and sections 3 and 4** Secondly, we need to define *p_N_(t)*, the probability of tumor survival based on highly fit mutant clones in the surrounding tissue. Following the assumptions in **sections 5.4 and 5.5**, we assume that there is a single mutant population *N* capable of removing tumors and that *N* mutant clones remove tumors as soon as they occupy the location of the tumor in the tissue. Our starting time occurs after the end of DEN treatment, and so much of the tissue may already be occupied by the *N* clones. Therefore we modify Equation 3 to account for the sensitive tumors existing in the remaining non-N proportion of the tissue.

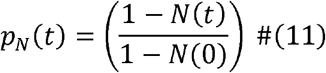

This still leaves us having to define *N*(*t*), the growth pattern of the mutant clones. Growth of mutant clones in EE have previously been modelled using branching processes, but these don’t consider competition between clones and limitations of the tissue size ^11,41^. Cellular automaton simulations have also been used to model clones in EE ^11^, but this does not allow for easy integration with the mathematical formulation. Instead, we used the logistic equation, which captures the key features of clonal spread: fast growth at early time points when mutant cells are mostly competing with surrounding wild type cells, slower growth at late time points when the tissue is already largely mutant, and an upper bound on total mutant spread ^11^. Therefore,

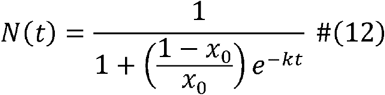

where *x*_0_ is the initial proportion of tissue covered by *N* mutant clones and *k* is the clone growth rate. To represent the experiment in which the DBZ Notch inhibitor is applied between 10 and 24 days after DEN, we can define

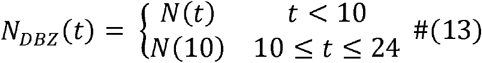

The full model can then be constructed by substituting these expressions into equation 9.

We have 7 independent parameters: *d*, Δ, *n*, *x*_0_, *k*, *S*(0), and the initial tumor density (**Supplementary information Table A**). We used Approximate Bayesian Computation (ABC) to find the parameter combinations for which the model most closely matched the mean tumor density from 10 days to 18 months after DEN treatment and the mean tumor density in CTL and DBZ experiments at 24 days after DEN treatment (see **methods**). The parameters were constrained as listed in **Supplementary information Table A**. The results are shown in were (95% credible interval lower bound, upper bound) *d*=0.29/day (0.17, 0.35), Δ=0.020 **Supplementary information Figures D and E**. The median acceptable parameters found (0.007, 0.047), *n*=1.4 (1.0, 2.5), *x*_0_=0.80 (0.42, 0.99), *k*=0.032 (0.019, 0.040), *S(0)*=0.64 (0.44, 0.82) and initial tumor density=4.2/mm^2^ (3.4, 5.4), although, given the simplifications and approximations used in this numerical example of the model, they should not be interpreted as estimates of their biological counterparts.

**Supplementary information Figures D and E.**
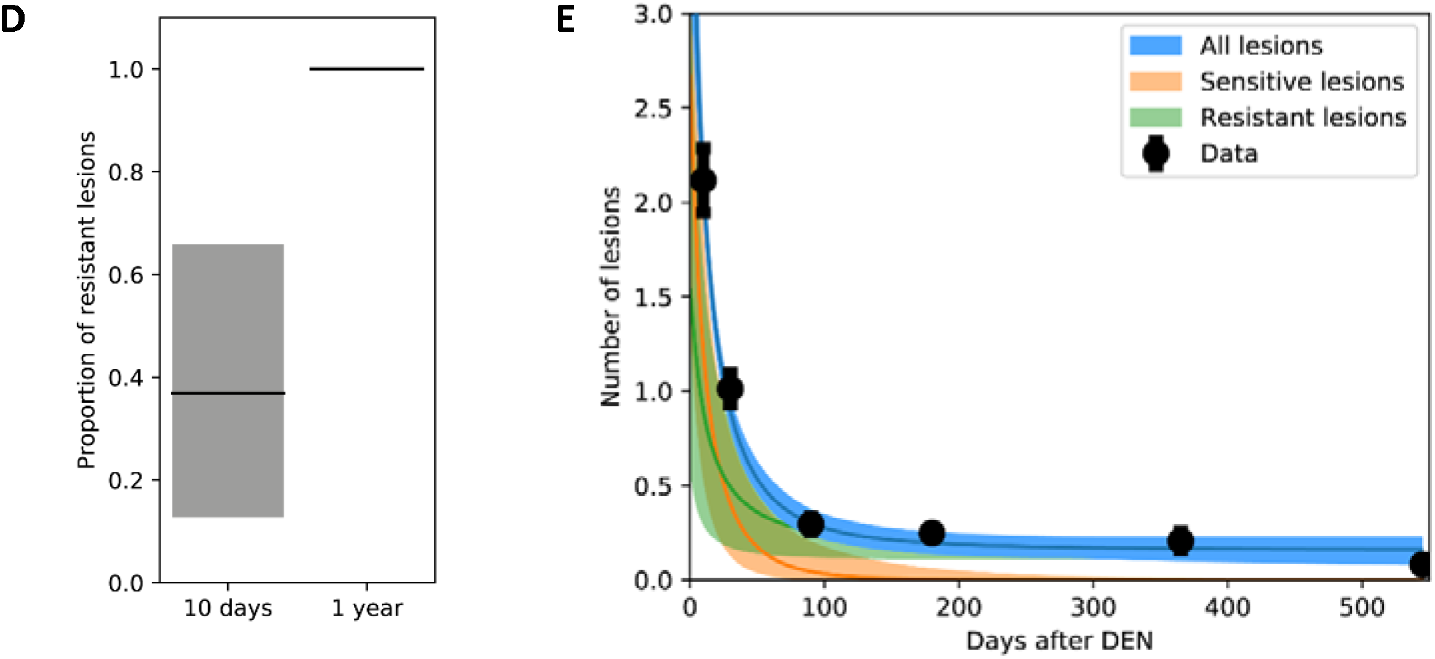
Numerical example of the model showing increased proportion of tumors resistant to displacement by mutant clones (**D**) and the decrease in lesion density (# tumors per mm of EE) following DEN-treatment (**E**). Model fit to tumor density data using Approximate Bayesian Computation (**supplementary information** and **Methods**). The mean value and range between the minimum and maximum outputs of the model run with the accepted parameters are shown.

The numerical example of the modelling principles demonstrates how the elimination of (a subset of) tumors by mutant clones in the surrounding tissue leads to the experimental observations of decreasing tumor numbers following DEN treatment, higher tumor survival when competitive imbalance is removed (DBZ experiment), and a selection pressure on tumor genotype.

##### 5.7 Summary

Together, the experimental data and modelling indicates that tumors in the DEN-treated EE are eliminated by mutant clones in the surrounding normal tissue. The data also suggests that the tumor genotype influences the chance of a tumor surviving, possibly by allowing the tumors to better compete with mutant clones in the surrounding tissue.

**Supplementary text Table A.**
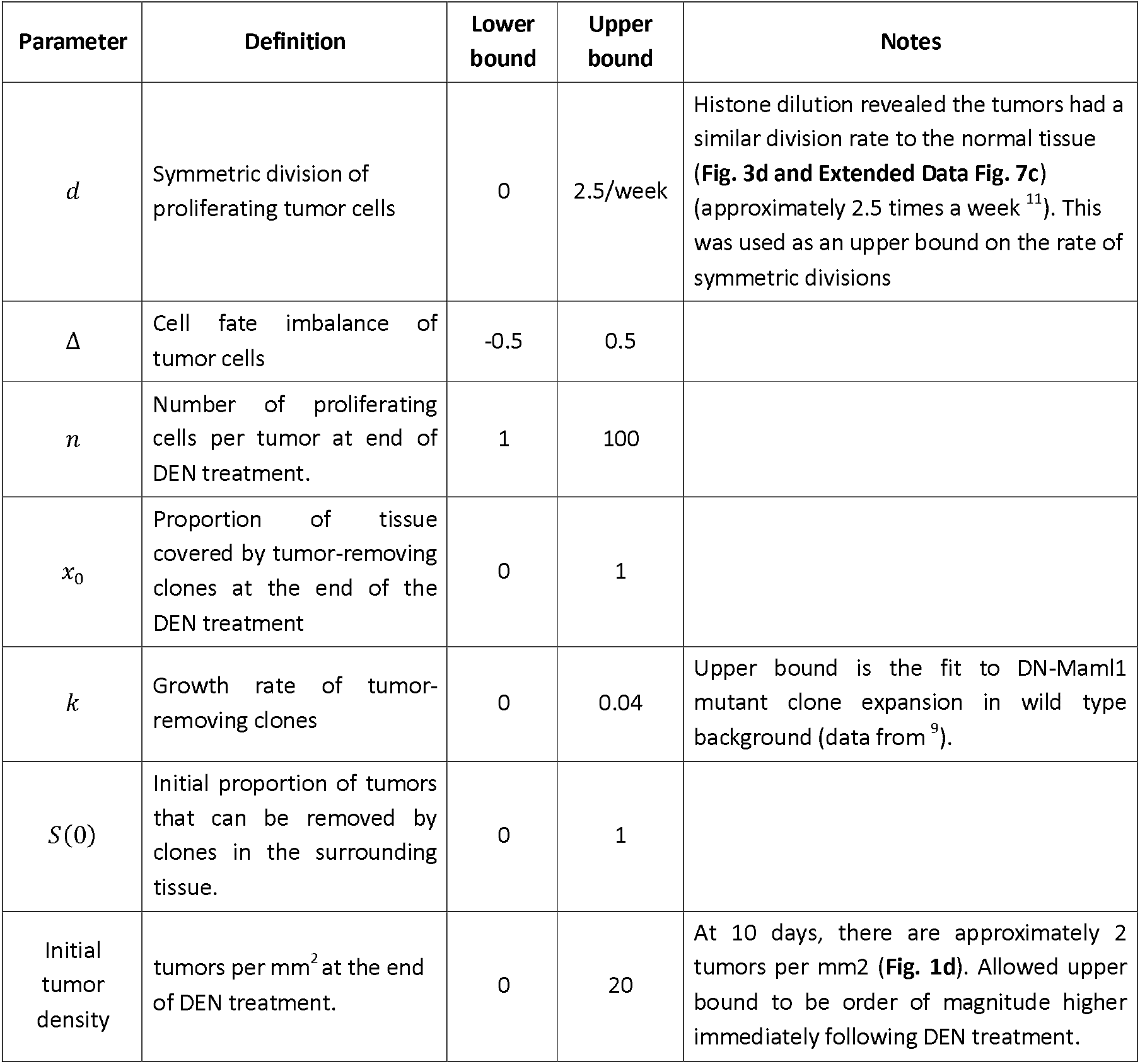

---

**Extended Data Fig. 1.**
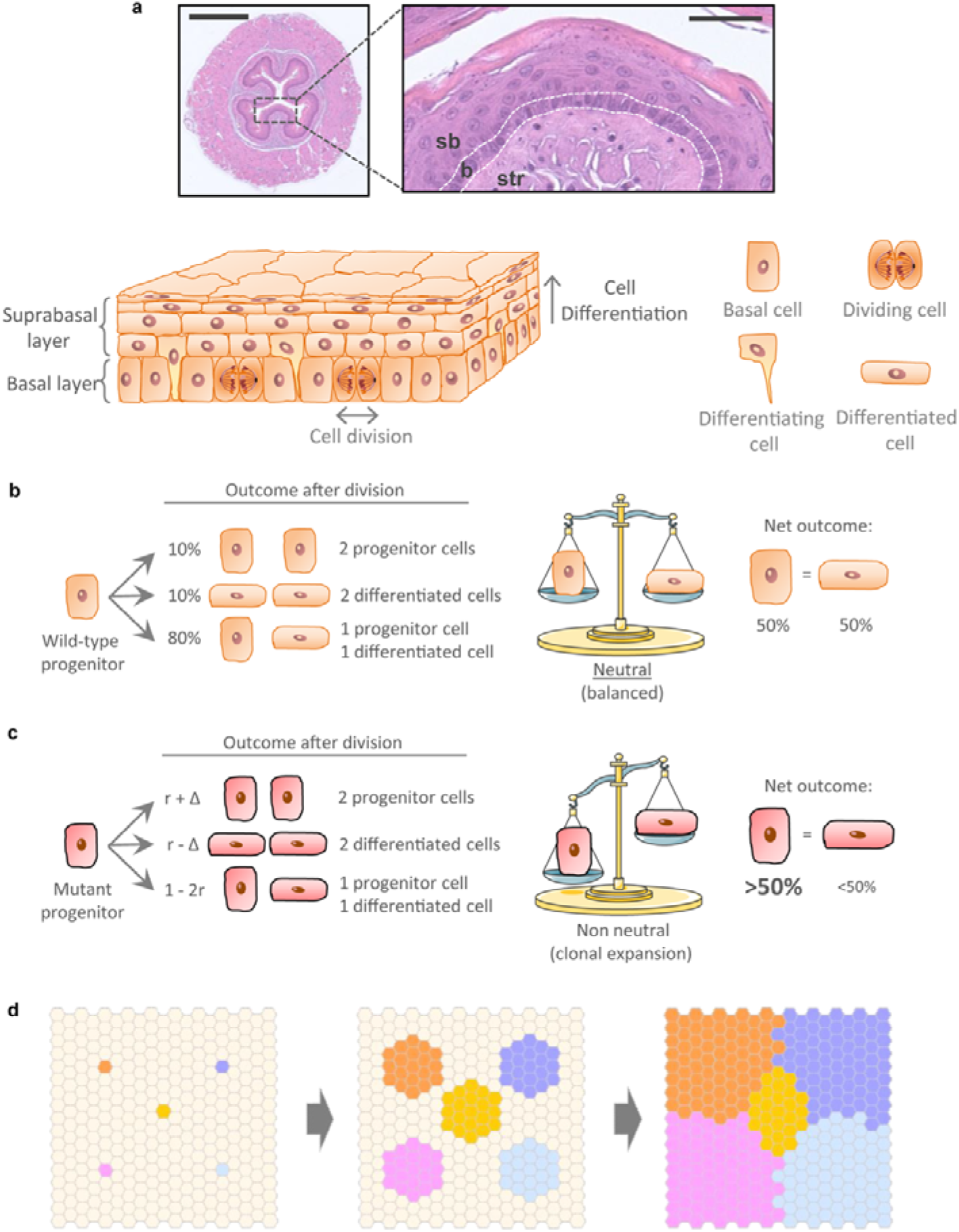
Mouse EE architecture and dynamics. (a) H&E images and schematic of the mouse EE architecture and dynamics; b = basal cell layer (delineated by white dots), sb = suprabasal layers, str = stroma. Scale-bars: 500μm (top), 50μm (bottom). Dividing (progenitor) cells are confined to the basal layer. Differentiating cells exit the cell cycle, migrate out of the basal layer, through the suprabasal layers, and are finally shed. (b-c) The single progenitor model (Supplementary information). All progenitor cells in the EE basal layer are functionally equivalent and following division produce either: two progenitors that will persist in the tissue, two differentiating cells that will cease division, stratify and be lost or one cell of each type. In homeostasis (b), the likelihood of the two progenitor and two differentiating cell outcomes is equal. Mutations (c) may tip the balance towards a non-neutral behavior, resulting in clonal growth if they favor proliferation of daughter cells. (d) Expansion of mutant clones is defined by their relative competitive fitness to adjacent clones. Initially, a fit “winner” mutant progenitor (colored) shows a fate bias towards proliferation and outcompetes its less fit “loser” surrounding cells, resulting in clonal expansion. Eventually, mutant clones begin to collide with each other until surrounded by similarly competitive mutants, at which point their cell fate reverts towards balance and their expansion slows (Supplementary information).

**Extended Data Fig. 2.**
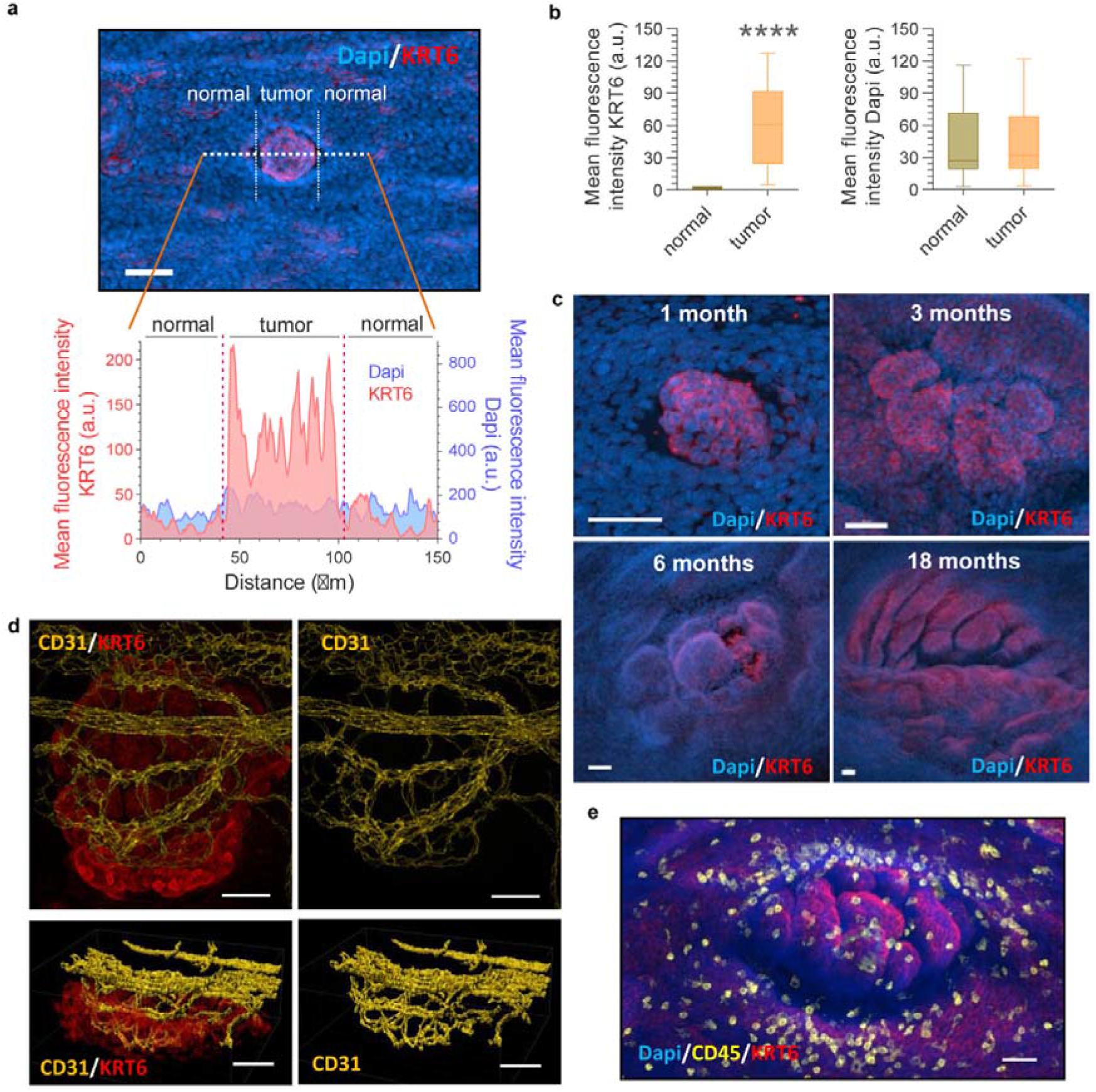
Characterization of DEN-generated tumors. (a) Image of a tumor (10-day post-DEN) stained with KRT6 and Dapi. Graph show the mean fluorescence intensity of KRT6 and Dapi measured across the white dotted line in the image above, illustrating the expression levels in the tumor and the adjacent normal epithelium. (b) Quantification of the mean fluorescence intensity of KRT6 and Dapi in tumors and their normal adjacent epithelium (as in a). ****P<0.0001 (Two-sided Mann-Whitney test, n=48 tumors from 3 mice). (c) Confocal images of mouse EE tumors collected at the indicated time points post-DEN treatment. (d) Confocal image of a 3-month post-DEN tumor stained for the endothelial cell marker CD31. Top and bottom images show top and lateral projections, respectively. (e) Confocal image of a 3-month post-DEN tumor stained for the immune cell pan-marker CD45. Scale-bars: 50μm.

**Extended Data Fig. 3.**
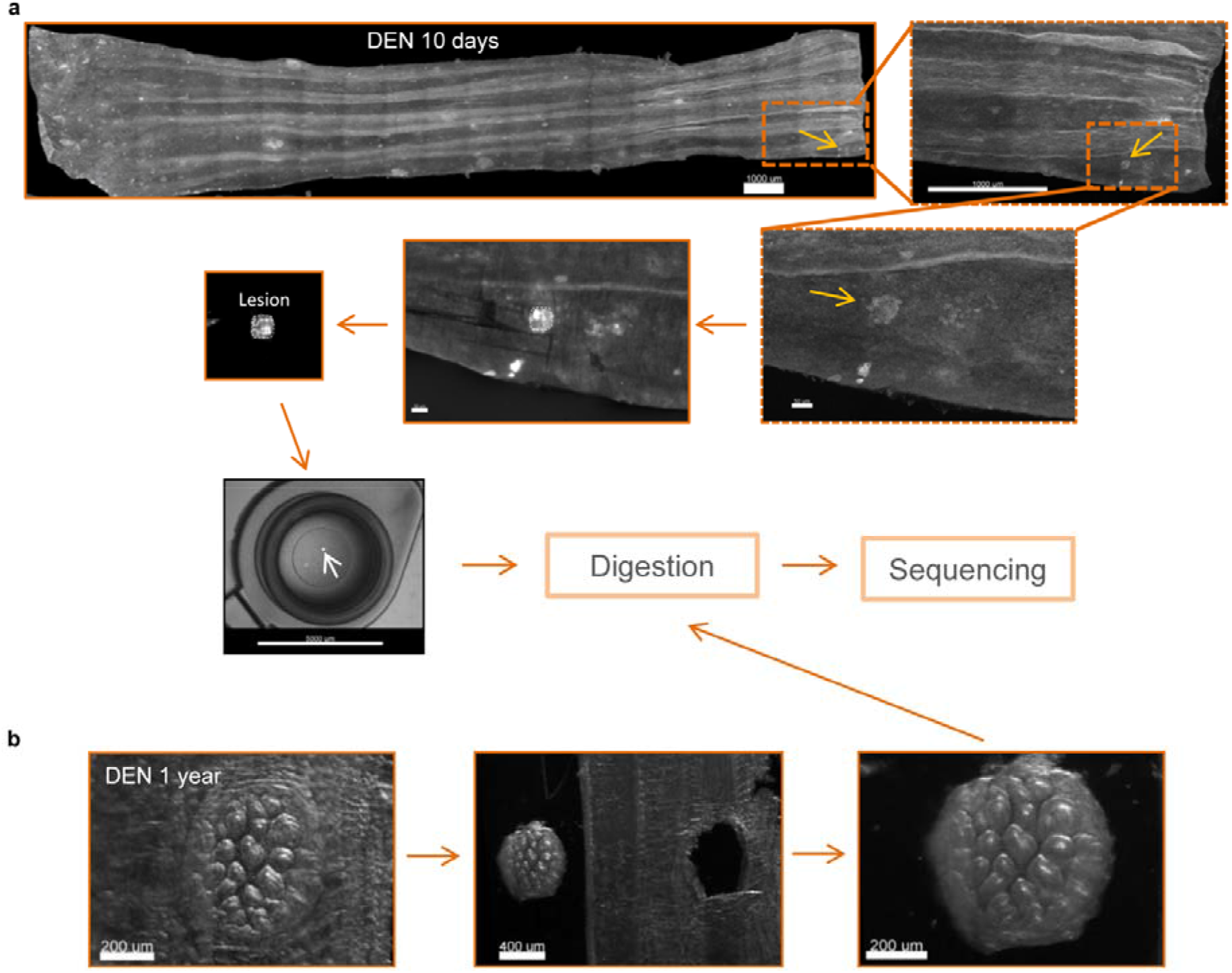
Collection of 10-day and 1-year EE tumors. (a-b) Esophagus were collected 10-days (a) or 1-year (b) post-DEN treatment. Tissues were longitudinally cut opened and the epithelium separated from the underlying muscle and stroma. Whole EEs were flattened, fixed, stained with KRT6 (grey), mounted and 3D-imaged on a confocal microscope. Tumors were identified from the processed images and manually cut under a fluorescent microscope. Samples were placed on 0.2ml tubes, digested and sequenced as detailed in Methods.

**Extended Data Fig. 4.**
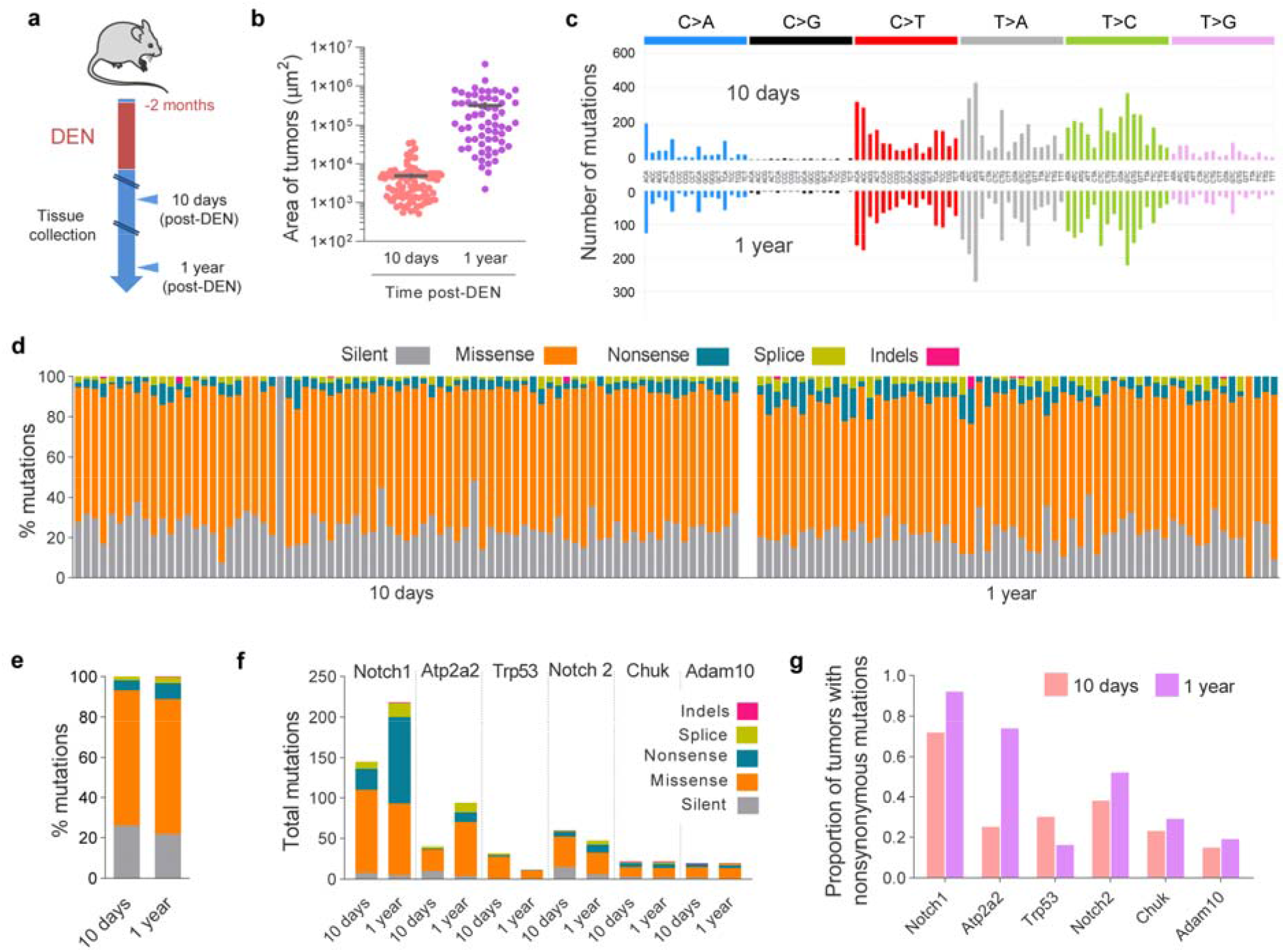
Targeted sequencing of 10 days and 1 year tumors. (a) Protocol: wild-type mice were treated for two months with DEN and the EE tumors collected 10-days or 1-year later. Samples were sequenced with a targeted approach (192 gene panel). (b) Area of 10-day and 1-year post-DEN tumors. Lines show Mean ± s.e.m. (n=89 and 64 tumors, respectively). (c) Mutational spectrum of 10-day and 1-year tumors. The bar plots illustrate the number of mutations in each of the 96 possible trinucleotides. (d-e) Percentage of silent, missense, nonsense and splice mutations and indels identified in 10-day or 1-year tumors. Graphs show the values for individual tumors (d, each column is a tumor), or the average for all tumors at each time point (e). (f) Number and type of mutations in the positively selected genes identified by dN/dS analysis from 10-day or 1-year tumors. (g) Proportion of 10-day and 1-year tumors carrying nonsynonymous mutations in the positively selected genes.

**Extended Data Fig. 5.**
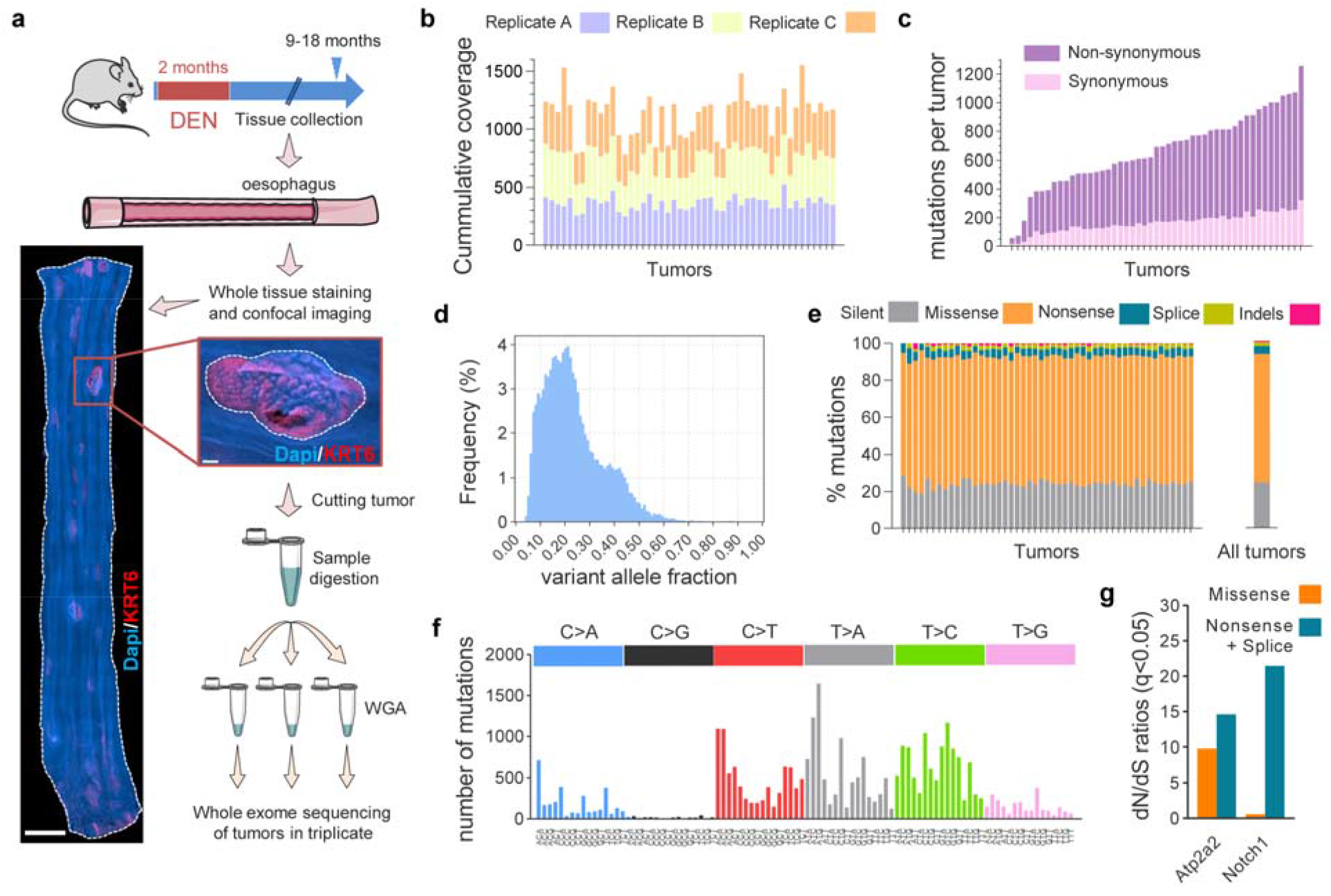
Whole exome sequencing of surviving tumors. (a) Wild-type mice received DEN for two months and the esophagus was collected 9-18 months after treatment. Tissues in (b) were stained for Dapi (blue) and KRT6 (red) and confocal imaged to identify tumors. Scale-bars = 2mm (main), 150μm (inset). Individual EE tumors were manually cut under a fluorescent microscope, digested and separated in triplicates. Each triplicate was whole genome amplified (WGA) followed by whole exome sequencing. To exclude artefactual SNVs generated during WGA, only mutations shared by all three amplified triplicates were considered for further analysis. A total of 32,736 mutations, including silent, missense, nonsense and splice mutations and indels were identified from 49 tumors in 16 mice. (b) Cumulative sequencing coverage of the whole exome triplicate samples. (c) Number of synonymous (light colored) and non-synonymous (dark colored) mutations per tumor, ranked by mutation burden. (d) Distribution of the variant allele fraction (VAF) for the mutations common within triplicates in each tumor. (e) Percentage of silent, missense, nonsense and splice mutations and indels for individual tumors and for all tumors combined. (g) Mutational spectrum of the tumors. The bar plots illustrate the percentage of mutations in each of the 96 possible trinucleotides. (g) dN/dS ratios for missense and truncating (nonsense + essential splice site) substitutions indicating genes under significant positive selection in the tumors (q<0.05, calculated with R package dNdScv).

**Extended Data Fig. 6.**
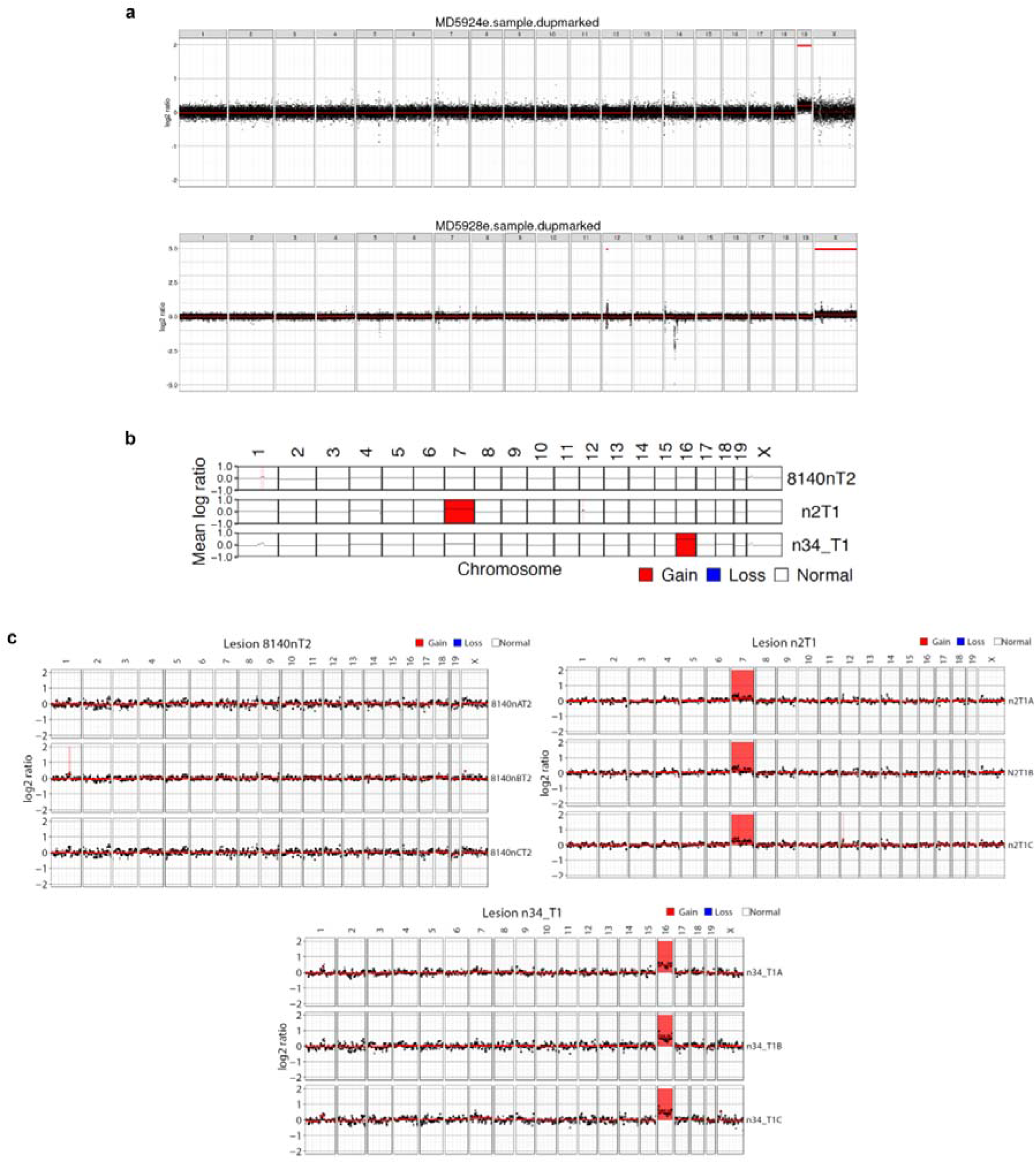
Analysis of chromosomal alterations in 1-year tumors. (a) Analysis of CNAs by whole genome sequencing of 1 year post-DEN tumors. Only 2 out of 64 tumors, MD5924e and MD5928e, exhibited CNAs. (b) Summary of chromosomal alterations found by whole exome sequencing data of 1-year post-DEN tumors in triplicate. 8140nT2 (top) is a representative example of a tumor without chromosomal alterations. 2 out of 49 tumors (n2T1 and n34_T1) showed small alterations (indicated by color). Only alterations present in all 3 triplicates (c) were considered valid calls.

**Extended Data Fig. 7.**
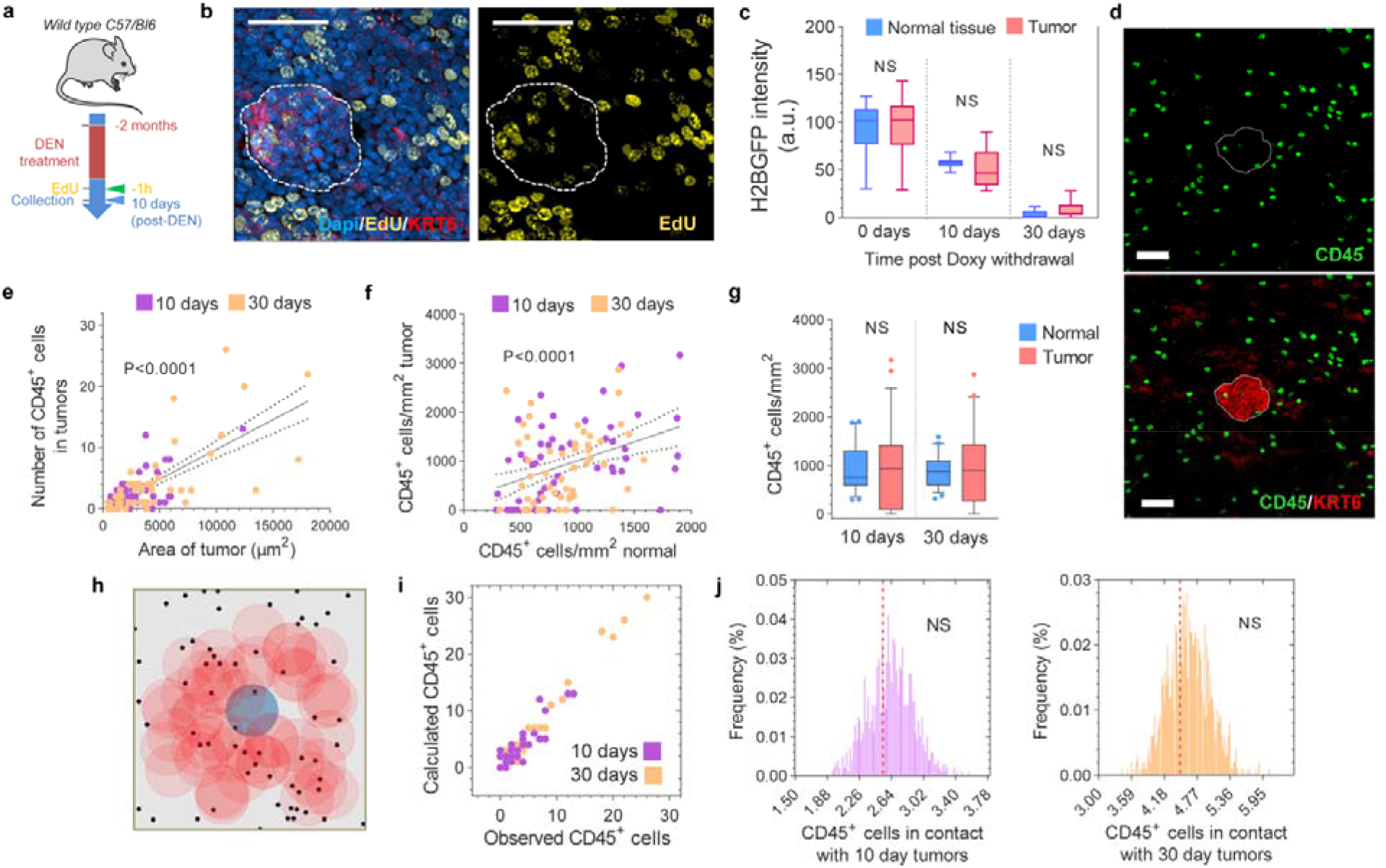
Early neoplasms are not eliminated by tumor cell abnormal proliferation or the immune system. (a) Protocol: wild-type mice were treated with DEN for two months and the tissues collected ten days later. Mice received EdU 1h before tissue collection. (b) Confocal images showing EdU incorporation (1h pulse) into tumors (dotted line) and the surrounding normal epithelium (n=22 tumors, 3 mice). Scale-bars: 20μm. See Supplementary information. (c) HGFP intensity in tumors and normal EE at the indicated time points post-Doxy withdrawal. (Two-sided Mann-Whitney test, n= 10 images per group). (d) Confocal image depicting CD45^+^ (immune) cells (green) within a 10-days post-DEN tumor (white dotted line) and its adjacent normal epithelium. (e-f) Correlation between the size of the tumors and the number of CD45^+^ cells within them (e) and the number of CD45^+^ cells in the tumor and the normal epithelium (f). Lines shows the Pearson correlation: R^2^ =0.5039 (e), and R^2^ =0.1464 (f), with 95% confidence interval (dotted lines). (g) Quantification of the number of CD45^+^ cells per mm^2^ of tumor and normal EE from wild-type mice 10- or 30-days post-DEN treatment (Two tailed Wilcoxon matched-pairs test, n= 53 and 50 images, respectively). (h) Permutation analysis of leukocyte location within tumors, based on the experimental measurements obtained from (d). For each image, the location of CD45^+^ cells (black dots) was left intact while the location of the tumor (colored circles) was randomly shuffled (blue shows original location, red shows shuffled “phantom” tumors), and the number of immune cells in contact with the tumor was counted. This was repeated 1000 times to produce the expected distribution, assuming no association between tumor and immune cell locations. (i) Calculated (within the original location) vs. experimentally observed number of CD45^+^ cells in 10- or 30-days post-DEN tumors. (j) Distribution of the average number of CD45+ cells in 10- or 30-days post-DEN tumors, obtained from the permutation analysis in (h). Red dotted line shows the experimentally observed average number of CD45^+^ cells within tumors at the indicated time-point. See **Supplementary information**. Box-plots in (c) and (g) show median (central box line) 25^th^-75^th^ percentiles (box), 1^th^−99^th^ percentiles (whiskers) and outliers (dots). NS= not significant.

**Extended Data Fig. 8.**
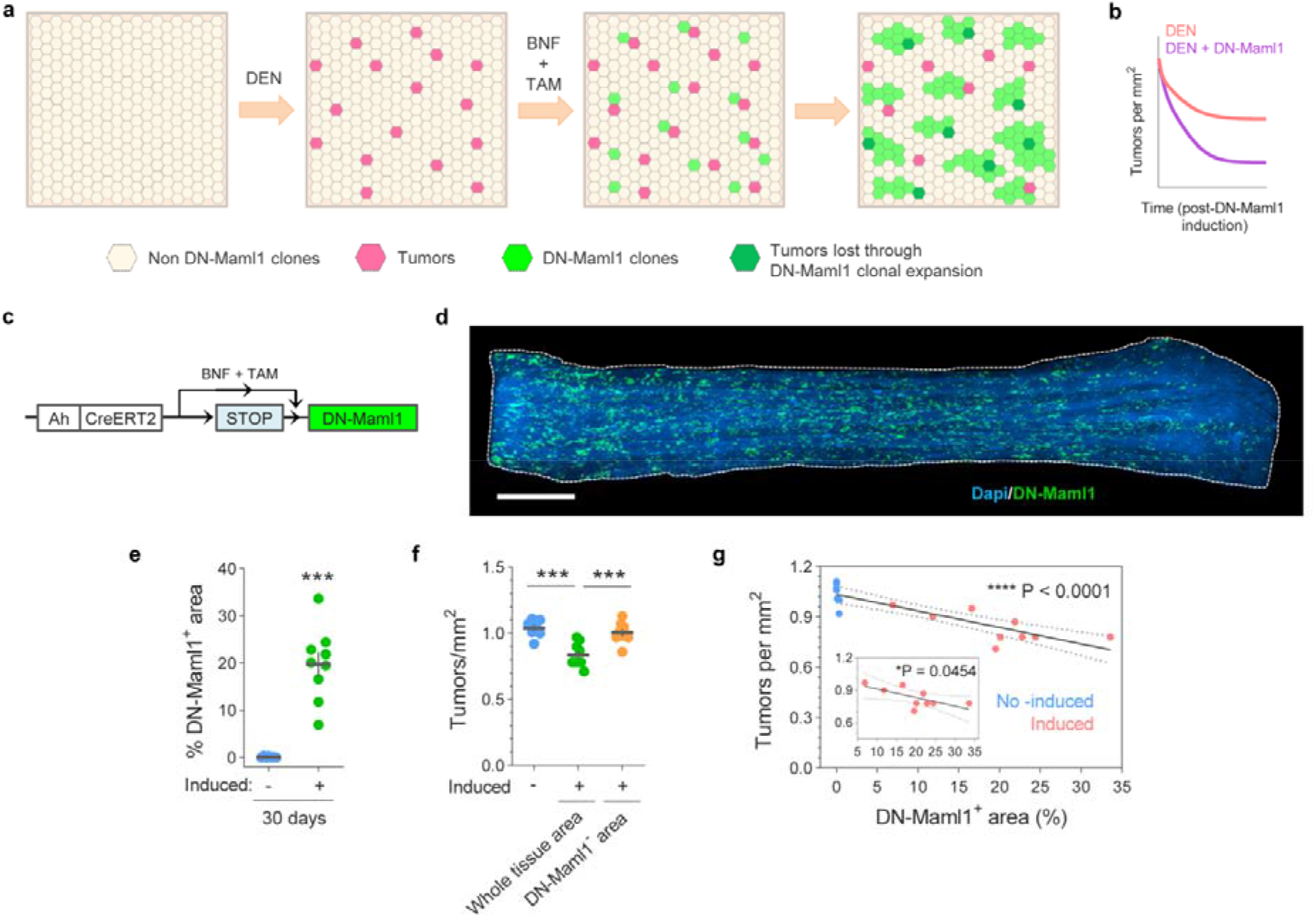
Expansion of highly competitive clones in normal tissue eliminates early tumors. (a) Cartoon illustrating the elimination of tumors due to competition with the highly competitive DN-Maml1 clones. Tumors are generated by DEN treatment followed by induction of DN-Maml1 clones with BNF and TAM. As DN-Maml1 clones expand they eliminate less fit tumors. (b) Qualitative representation of tumor dynamics following DN-Maml1 induction as compared to non-induced controls (see **Supplementary information**). (c) *In vivo* genetic lineage tracing using *Ahcre^ERT^Rosa26^wt/DNM-GFP^* reporter mice. Upon injection of the drugs tamoxifen (TAM) and ß-napthoflavone (BNF), Cre-mediated recombination results in the heritable expression of the highly competitive dominant negative allele of Maml-1 fused to GFP fluorescent protein (DN-Maml1), which will then be expressed in all the progeny of the single marked cells, generating clusters of labelled mutant clones. (d) Confocal image of a DEN-treated *Ahcre^ERT^Rosa26^wt/DNM-GFP^* mouse whole EE, depicting DN-Maml1 clones (green). Mice were induced ten days after DEN withdrawal and esophagus collected twenty days later (as in Fig. 4b). Scale-bar = 2mm. (e) Percentage of EE area covered by DN-Maml1 clones in induced and non-induced (control) mice (n= 9 and 7 mice, respectively). Error-bars are Mean ± s.e.m. ***P<0.001 (Two-sided Mann-Whitney test). (f) Number of tumors per mm^2^ of EE in control non-induced (n= 7) and induced (n= 9) *Ahcre^ERT^Rosa26^wt/DNM-GFP^* mice. Data for the induced mice is show as the number of tumors per mm^2^ of whole EE (whole tissue area) and per mm^2^ of EE not covered by DN-Maml1 clones (DN-Maml1-area). Error-bars are Mean ± s.e.m. ***P<0.001 (Two-sided Mann-Whitney test). (g) Correlation between the area of EE covered by DN-Mam1 clones and the number of tumors in induced and non-induced *Ahcre^ERT^Rosa26^wt/DNM-GFP^* mice. Line shows the Pearson correlation (R =0.7526) with 95% confidence interval (dotted lines). Inset shows the correlation of the induced mice only (R =0.4577).

**Extended Data Fig. 9.**
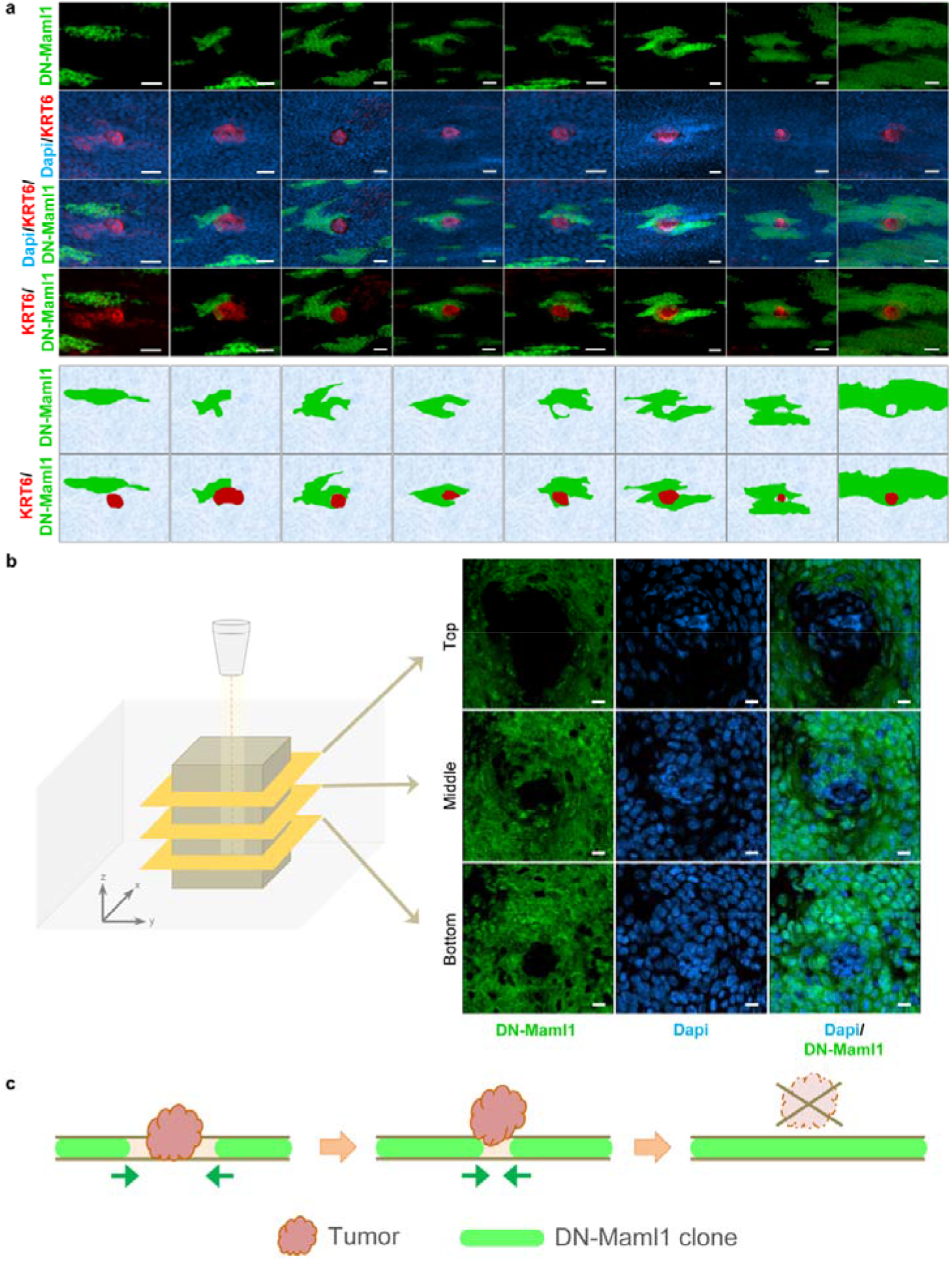
Spatial interaction between DN-Maml1 clones and tumors. (a) Confocal images and cartoons of EE from AhcreERTRosa26wt/DNM-GFP mice treated with DEN for 2 months, induced 10-days post-DEN withdrawal and tissues collected 20-days later. The images show different steps of DN-Maml1 clones (green) interacting with early tumors (red), from barely touching (left) to completely surrounding the tumors (right). Scale-bars: 25μm. (b) Top-down confocal images depicting a DN-Maml1 clone surrounding a tumor (see also Fig. 4e). Scale-bars: 10μm. A smaller tumor surface area at the base strongly suggest a process of extrusion by the expanding DN-Maml1 clone as illustrated in (c).

**Extended Data Fig. 10.**
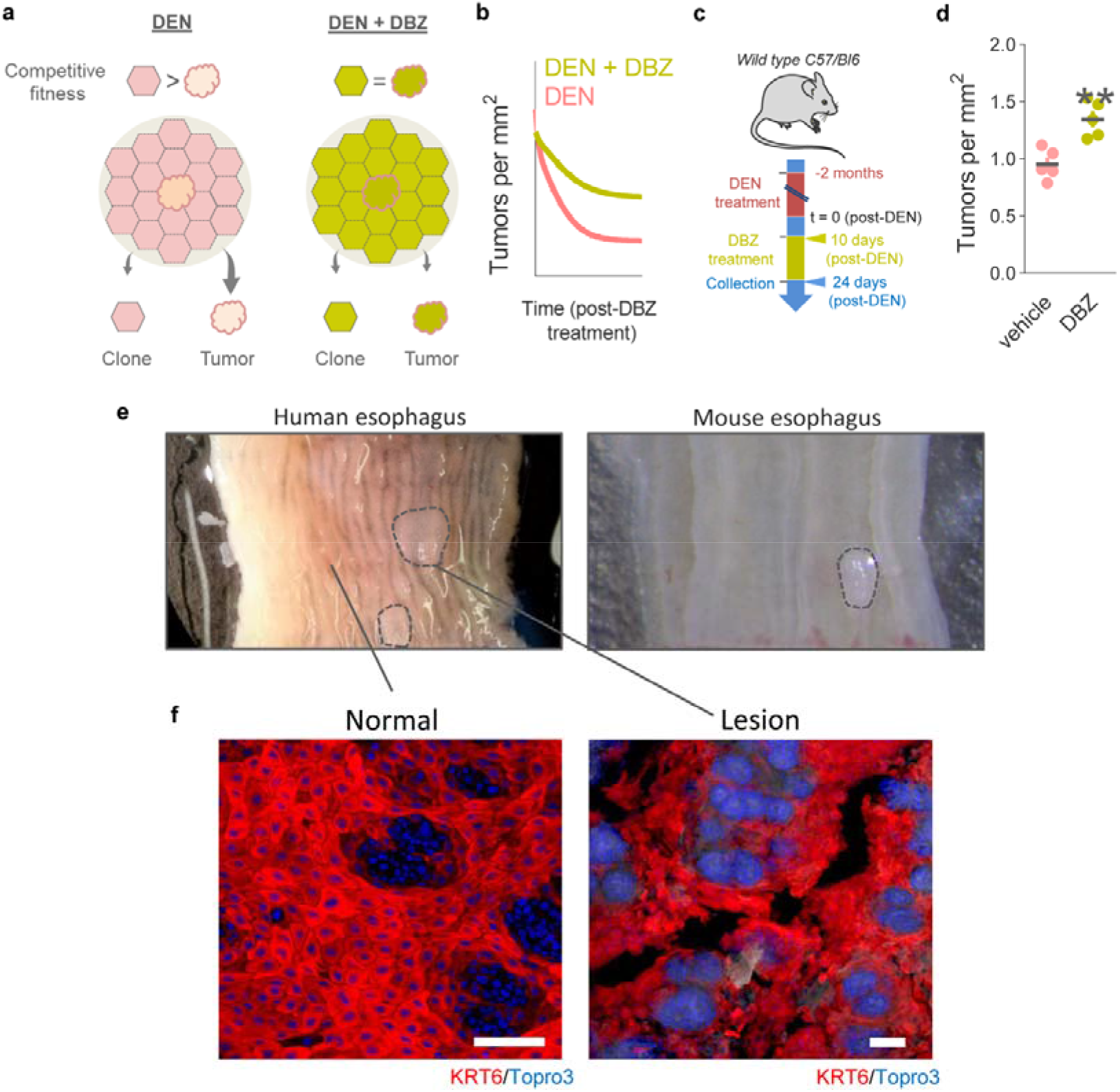
Increasing competitive fitness throughout EE slows down tumor loss. Human esophageal lesions. (a) Cartoon illustrating the predicted effects on tumor burden after leveling up the competitive fitness in normal and neoplastic EE through administration of DBZ. (b) Qualitative representation of tumor dynamics following DBZ treatment as compared to vehicle-treated controls. (c) Protocol: wild-type mice were treated with DEN for two months. Ten days after DEN withdrawal mice received DBZ or vehicle control and tissues were harvested two weeks later. (d) Number of tumors per mm^2^ of EE in DBZ and vehicle control treated mice. Error-bars are Mean ± s.e.m. **P=0.0079 (Two-sided Mann-Whitney test, n=5 mice per group). (e) Images of human (left) and mouse (right) esophagus. Tissues were longitudinally cut opened and flattened. Dotted lines delineate lesions. Scale-bars: 1mm. (f) Confocal images of normal (left) and neoplastic (right) epithelium stained with KRT6 (red) and Topro3 (nuclei, blue). Scale-bars: 100μm.

